# Multiparametric domain insertional profiling of Adeno-Associated Virus VP1

**DOI:** 10.1101/2023.04.19.537549

**Authors:** Mareike D. Hoffmann, Alina C. Zdechlik, Yungui He, David Nedrud, George Aslanidi, Wendy Gordon, Daniel Schmidt

## Abstract

Evolved properties of Adeno-Associated Virus (AAV), such as broad tropism and immunogenicity in humans, are barriers to AAV-based gene therapy. Previous efforts to re-engineer these properties have focused on variable regions near AAV’s 3-fold protrusions and capsid protein termini. To comprehensively survey AAV capsids for engineerable hotspots, we determined multiple AAV fitness phenotypes upon insertion of large, structured protein domains into the entire AAV-DJ capsid protein VP1. This is the largest and most comprehensive AAV domain insertion dataset to date. Our data revealed a surprising robustness of AAV capsids to accommodate large domain insertions. There was strong positional, domain-type, and fitness phenotype dependence of insertion permissibility, which clustered into correlated structural units that we could link to distinct roles in AAV assembly, stability, and infectivity. We also identified new engineerable hotspots of AAV that facilitate the covalent attachment of binding scaffolds, which may represent an alternative approach to re-direct AAV tropism.

## INTRODUCTION

Recombinant Adeno-associated virus (rAAV) has proven to be safe and able to drive long-term expression in dividing and non-dividing human cells. Several of AAV-based therapeutics have been approved by the FDA (1–4) and numerous clinical trials using AAV for the treatment of genetic diseases are underway. Despite the exceptional clinical potential of naturally evolved AAV serotypes, they could be substantially improved with respect to production yield, DNA packaging capacity, immunogenicity, cell type specificity, and infectivity (reviewed in (5)).

Addressing these drawbacks is facilitated by the relatively simple structural and genetic organization of AAV. The ∼4.7kb single-stranded DNA genome of AAV comprises two genes, *rep* and *cap*, flanked by inverted terminal repeats (ITR). One ORF of the capsid gene *cap* encodes for three viral proteins VP1 (737 aa, 87 kDa), VP2 (600 aa, 72 kDa), and VP3 (535-503 aa, 62 kDa). It is expressed from the p40-promoter (C-terminus of the *rep* gene) and translated from overlapping open reading frames in a way that VP2 is lacking the N-terminus of VP1, and VP3 is missing the N-terminal part of VP1/VP2 (6). Other *cap* ORFs express the assembly-activating protein (AAP) (7) and membrane-associated accessory protein (MAAP (8)). 60 VP monomers assemble into a viral capsid at an average ratio of 1:1:10 VP1, VP2, and VP3, respectively (9–12). This ratio is highly divergent and capsid assembly is stochastic, such that every capsid has a unique structural assembly (13). The icosahedral capsid features a cylindrical pore at the 5-fold interface, depressions surrounding the 5-fold pore continuing through the 2-fold axis, as well as protrusions at the 3-fold axis (14). Although the overall topology of AAV capsids is conserved across serotypes, Govindasamy et al. determined variable regions (VR1-9) mapping to surface loops of the capsid (15). The unique N-terminal portion of VP1, also known as VP1u, is located inside the capsid and is indispensable for infection. Upon infection, acidification during endosomal trafficking causes unfolding of the VP1u domain so that it can be externalized through the pore (16–20). A conserved phospholipase A2 domain (PLA2 (21)) and nuclear localization signals (NLS (22)), which are part of VP1u, can then facilitate endosomal escape and nuclear entry (23).

A variety of capsid engineering approaches have been applied in the past to improve the natural infection efficiency of AAVs ranging from shuffling of natural AAV serotypes, recovery of ancestral serotypes, and peptide display. These methods resulted in significantly advanced capsid variants, such as AAV-DJ (24), Anc80 (25), AAV-PHP.eB (26) or AAV2.7m8 (27). Moreover, there have been a few attempts to incorporate larger, structured protein domains. The first was the fusion of a green fluorescent protein (GFP) N-terminally to VP2, which was used to visualize intracellular trafficking of AAV particles (28). In the same manner, *Gaussia* Luciferase (29) and an ankyrin repeat protein (DARPin (30)) were successfully incorporated into the capsid. Other studies incorporated domains for a cell type-specific targeting into VR4 of VP1 or VP2, including nanobodies (31), HUH tags (32) or DARPins (33). While the aforementioned domain insertions were either N-terminal fusions or insertions into VR4 and peptide insertions predominantly focused on VR8, few studies attempted to comprehensively survey permissive capsid regions for domain insertions. Judd et al. constructed a random insertion library of mCherry into the VP3 encoding section of VP1 of AAV2, identifying only a single clone, in VR4, that tolerated insertion (34). Thus, there is a paucity of large-scale domain insertional datasets that comprehensively assess (i) the effect on biologically relevant functions, i.e., packaging, binding, and infection, as well as (ii) the effect of inserting domains with different physicochemical properties. In absence of these data, the boundaries of AAV capsid plasticity with respect to accepting domain insertions while maintaining fitness (i.e., assembly, packaging, cell entry, etc.) are yet to be fully understood.

Here, we combined Saturated Programmable Insertion Engineering (SPINE (35)) with sequencing-based fitness assays to comprehensively determine multiple AAV fitness phenotypes upon insertion of the FLAG peptide tag as well as several large, structured protein domains into VP1 of AAV-DJ. Our data revealed a surprising robustness of AAV viral capsids to accommodate large protein domain insertions. We also found strong positional, domain-type, and fitness phenotype dependence of insertion permissibility, which we can map to contiguous structural units of the AAV capsid and link to distinct roles in AAV assembly, stability, and infectivity. We also identified new engineerable hotspots of AAV accepting insertion of small protein tags that facilitate the covalent attachment of antibodies. These hotspots may enable alternative approaches to redirect AAV tropism.

## RESULTS

In the context of recombinant AAV as a gene therapy vector, capsid structural features impinge on a multitude of distinct virion functions. First, AAV must be produced recombinantly, which entails a stochastic oligomerization process forming an empty capsid in the nucleus that is governed by ordered interaction between capsid protein monomers (13, 36, 37), followed by packaging of a single-strand DNA payload through the 5-fold pore that is braced by complex subunit interactions (16, 38–40). The resulting virion must be sufficiently stable to survive biochemical purification after lysis of the producer cells (41).

Next, virions must attach to (co-)receptors on the surface of a target cell (AAV-DJ: heparan sulfate (42, 43)), followed by endocytosis (23). After endocytosis, virions must interact with the recently identified universal AAV receptor (AAVR) to ensure proper trafficking to the trans-Golgi network (23, 44, 45), undergo the required conformational changes (autoproteolysis, externalization of the N-terminal PLA2 domain (19, 21, 46)) that together mediate virion escape into the cytosol. Once in the cytosol, several intracellular trafficking events remain: capsids interact with the nuclear pore complex, enter the nucleus, and are forwarded to the nucleolus, where the genetic payload is released (20, 23).

Our goal in this study was to survey as many proxies (‘AAV fitness phenotypes’) for these distinct virion functions as possible. We posit that the resulting comprehensive multiparametric AAV fitness datasets can facilitate the optimization of engineering AAV along multiple axes (8, 47, 48). Furthermore, based on our previous studies in ion channels (49, 50), we hypothesized that systematic domain insertion (i.e., perturbation scanning) across different measured phenotypes may uncover the underlying topological organization of AAV capsids and provide insight into capsid determinants for assembly, stability, and dynamics of virus capsids.

### Domain insertional profiling in AAV-DJ VP1

We turned to AAV-DJ (24) as the testbed for inserting a peptide tag (FLAG) and six different protein domains (Supplemental Figure 1) in between every two residues of VP1. Our rationale for focusing on VP1, as opposed to the more abundant VP3 or the non-essential VP2, was as follows: As the least abundant VP isoform there are, on average, between 1-5 copies of VP1 incorporated per capsid (9–12). Note that this is an average copy number based on bulk measurements (9–12); because assembly is a stochastic process, there are many particles with copy numbers at the extreme tails (i.e., 0 copies or > 10 copies) (13). We reasoned that keeping a low number of VPs carrying inserted domains, which are potentially very disruptive, is more likely to result in assembled and functional virions. This is akin to applying a low or intermediate amount of selection pressure in directed protein evolution experiments, which can reveal more facetted fitness phenotypes (51). Furthermore, we had observed in a prior study that inserting large domains into VR4 (part of the VP common region) completely abolished AAV production unless it was limited to VP1 (or VP2) only (32). Most importantly, focusing on VP1 allows us to interrogate the effect of domain insertion on AAV cell entry; VP1 is required for AAV infectivity as it contains the PLA2 domain that mediates endosomal escape (19, 21, 46).

To generate separate expression constructs for the VP1 domain insertion library and *cap* expressing only VP2 & VP3, we duplicated the AAV-DJ *cap* gene and introduced mutations (M1K; T138A/M203K/M211L/M235L) to suppress expression of VP1 or VP2/VP3, respectively (Figure 1A, Supplemental Table 1). We also replaced the VP1 heparin binding domain (HBD: residues 587-590) with an HA tag. The VP1-only *cap* gene was then subjected to SPINE (35), which resulted in a library of VP1 variants with a peptide tag or protein domain, flanked by short linkers, inserted in between every two residues. One motif we inserted was the FLAG peptide tag (DYKDDDDK), because of its similar size to prior AAV peptide insertions (52–54). With an eye towards redirecting AAV tropism, we focused on protein domains that themselves have retargeting abilities (nanobody binding to GFP (55)) or that enable covalent linkage of retargeting moieties (SNAP-tag to link O6-benzylguanine derivatives (56); SpyCatcher to link SpyTag fusions (57)), and three different HUH tags (WDV, DCV, mMobA) to covalently link ssDNA-conjugated molecules in a sequence specific manner (58–61).

**Figure 1:**
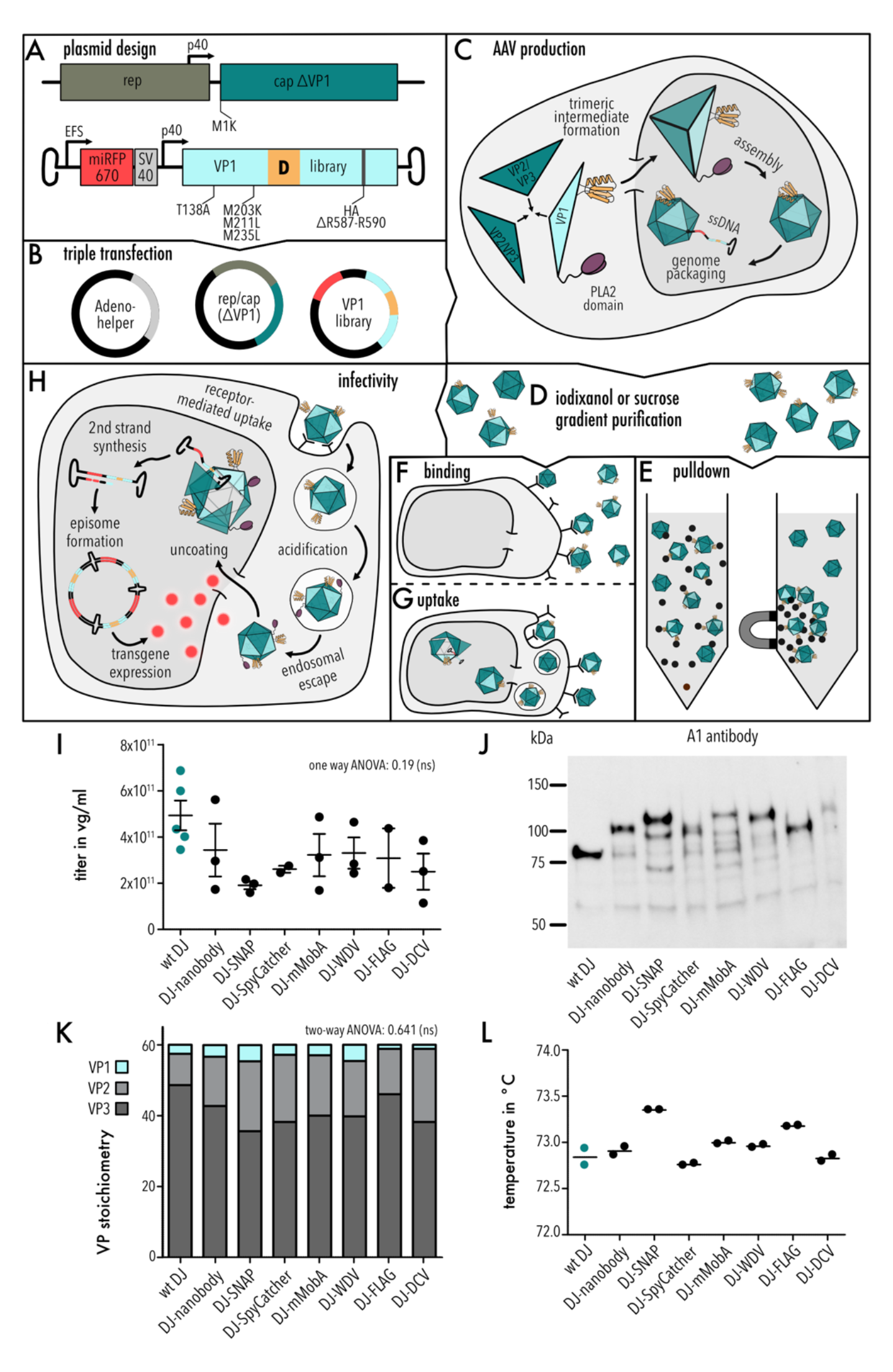
**Design and analysis of AAV domain insertion libraries.** (A) Schematic of library design. (B) Plasmids used for the triple transfection in AAV production. (C) Schematic of AAV capsid assembly. (D) Purification of AAV via gradient purification. (E-H) Schematic of workflows analyzing the fitness of AAV domain insertion libraries, i.e., pulldown, binding, uptake, and infectivity assays. (I) Quantification of packaging titer via qPCR. Data are means ± SEM. One way ANOVA test p value 0.19, not significant (ns). (J) Representative Western blot image of AAV domain insertion libraries stained with A1 antibody (detecting VP1 subunits). (K) Western blot quantification of VP1, VP2 and VP3 subunits. Data are means (n=3). Two-way ANOVA test p value 0.641, not significant (ns). (L) T_m_ of AAV libraries obtained by DSF assay. Data are means (n=2).

Since insertions can affect any of the steps in AAV packaging and infection, we attempted to independently assay different insertion variant fitness phenotypes. The relatively large insertion variant library size (744 AAV positions (including HA tag) x 7 motifs = 5,208 variants for each assay) necessitated a high throughput format. We therefore devised assays in which the different fitness phenotypes of an insertion variant were assessed by NextGen sequencing variant populations before and after a fitness test (62). A requirement for our approach was a stringent linkage between genotype (the insertion variant) and the measured phenotype (determined by properties of the capsid into which this VP1-variant is assembled). We achieved this by flanking the VP1 variant library with ITRs such that the gene encoding a specific VP1 variant became packaged into the capsid that incorporated this variant during assembly. We avoided the potential pitfall of cross-packaging (a mismatch between packaged VP1 variant gene and VP1 variant protein that incorporated into the capsid) by transfecting producer cells at very low MOI. This has been demonstrated to reduce cross-packaging (63).

We then used these VP1 variant input libraries, stratified by inserted motif, for helper-free virus production followed by gradient purification (Figure 1B-D). Using NextGen sequencing (NGS) of packaged genomes, we assayed ‘AAV packaging fitness’ by counting the frequency of a given VP1 variant (*i*) after packaging (*s*) relative to the frequency of that variant in the input library (*u*), normalized to wildtype AAV (*wt*):

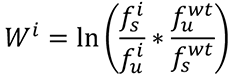

We measured absolute wildtype fitness by spiking in an AAV-DJ ITR flanked VP1 only *cap* gene containing 10 synonymous mutations into the VP1-library mix (Supplemental Table 1). Wildtype AAV-DJ and AAV-DJ with silent mutations showed no difference in production titers (Supplemental Figure 2). Using a similar approach to count VP1 variants before and after selection, we established assays that determine pulldown fitness (using affinity purification material), cell binding, cell uptake, and infectivity fitness (Figure 1E-H). Infectivity assays were based on transduction of HEK293FT cells with miRFP670nano (64) expressed from the AAV payload (Figure 1A). This enabled flow sorting of infectious variants (enriched in miRFP670nano^high^ cells) (Supplemental Figure 3).

### VP1 insertion library quality and completeness

Overall, coverage for all phenotype assays and inserted motifs was excellent (median dataset completeness is 98.5%; see Supplemental Table 2 for sequencing statistics). Biological replicates were highly correlated for plasmid library, packaging, and pulldown assays (Pearson correlation coefficient: 0.67-0.99, Supplemental Figure 4A). The infection assay was very noisy for some domains (Pearson correlation coefficient 0.38-0.88) likely related to small number of cells collected for the miRFP670nano^high^ cell pool (Supplemental Figure 3). For all tested motifs, the majority of missing insertion positions (no data in either replicate, for example positions 428-445) was missing in all phenotypes and the input library, suggesting that the dropout rate was related to library construction (Supplemental Figure 4B,C). Since replicate 2 had a higher read depth and completeness overall, we used this dataset for further analysis. Median read depth across tested motifs and phenotype assays was 635 reads per position (Supplemental Figure 4D, Supplemental Table 2).

### Domain insertions do no alter bulk properties

After helper-free production and iodixanol gradient purification, we measured virus titers by qPCR. While titers appeared somewhat lower for insertion libraries, this difference was not significant (Figure 1I, one-way ANOVA p-value 0.19). Western blot with the A1 antibody (which recognizes a VP1-unique epitope) confirmed that VP1 was incorporated into virions for all libraries (Figure 1J). We next used the B1 antibody densitometry to determine bulk VP1/VP2/VP3 ratios (Figure 1K, Supplemental Figure 5). We found that wildtype AAV-DJ virion contains on average three VP1 copies per 60-mer capsid, in line with prior studies (12, 13). There was no significant difference in VP composition for the different insertion libraries with respect to VP1 (two-way ANOVA p-value: 0.641). As another bulk characteristic, we measured capsid melting temperatures using differential scanning fluorimetry. While SNAP and FLAG libraries had slightly elevated melting points (Figure 1L, Supplemental Figure 6), all libraries were overall remarkably similar and within 1°C range of AAV-DJ, suggesting that domain insertions in VP1 did not significantly impact capsid stability. We next used negative stain electron microscopy to compare full/empty capsid ratios of wildtype AAV-DJ, SNAP-, and nanobody libraries and found that full particles comprised between 80-90% of all samples (Supplemental Figure 7).

Taken together, these data suggest that, on average (i.e., considering the entire insertion library), bulk properties are not altered by domain insertions.

### High resolution AAV fitness profiles across different phenotypes

We would expect that different insertion types at different sites do have different impact on AAV fitness, so we turned our attention to insertion motif-and region-specific differences. These data are summarized in Figure 2, showing a heatmap of insertion fitness of all seven motifs in all 744 VP1 positions, segregated by measured fitness phenotypes (packaging, pulldown, binding, uptake, and infectivity). Fitness values are mapped from magenta over white to green, corresponding to lower to higher than wildtype AAV-DJ fitness (white). Note that fitness measurements are derived from AAV capsid with a variable copy number of VP1 located at random faces, which will affect phenotype penetrance. Poisson errors were generally low (Supplemental Figure 8) and only slightly elevated at the junctions between the 14 different fragments used in the SPINE-mediated assembly of the AAV insertion library. Median error with respect to dynamic range of each fitness assay was (0.102 / 2) log units = 5.1%. Overall, there was a strong dependence on insertion position and type of inserted domain for most assays, with the notable exception of uptake and infectivity.

**Figure 2:**
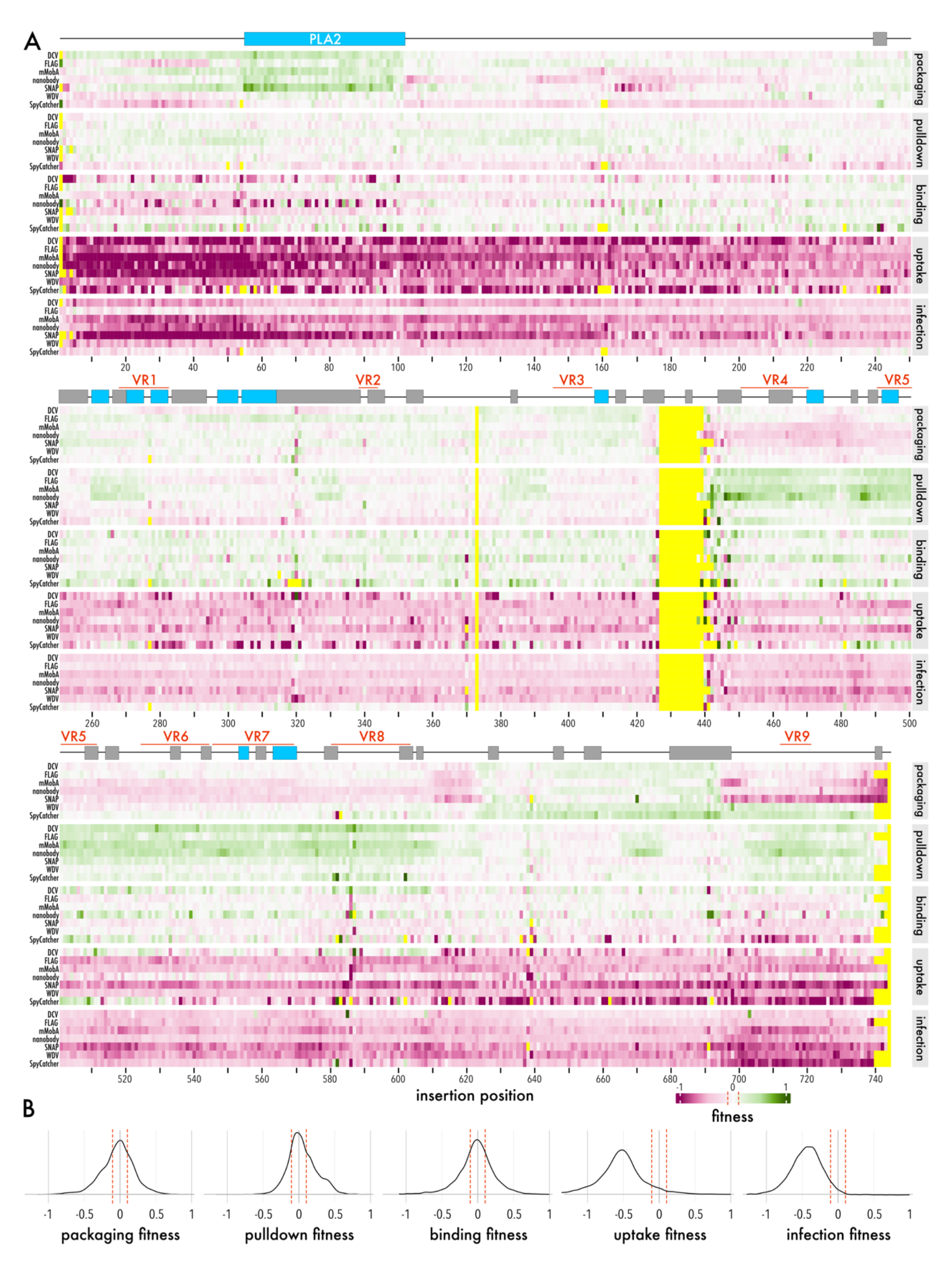
Fitness of AAV domain insertion libraries. (A) Heatmap representing the fitness of each motif insertion in each VP1 position for AAV packaging, pulldown, binding, uptake, and infectivity assays. Green indicates higher and magenta lower fitness than AAV-DJ (white). Yellow denotes positions without data. VP1 secondary structure elements and VR1-9 are indicated on top. (B) Fitness distributions of all AAV libraries compared to AAV-DJ (fitness = 0) ± standard error (red dashed lines).

### Effects of domain insertions in VP1u

Focusing on packaging fitness, we saw a distinct increase in fitness when motifs (in particular: nanobody, SNAP tag, or DCV HUH tag) were inserted into the PLA2 domain of VP1u (Figure 3). This suggests that perturbing the PLA2 domain at the assembly stage resulted in increased packaging efficiency. One possible explanation is that an insertion causes such a dramatic conformational change that VP1 becomes incompatible with trimer or pentamer formation, the initial steps of capsid assembly (36, 37). Malformed VP1 may ‘poison’ trimer assembly in the cytosol, may interfere with transport to the nucleus, or may interfere with trimer addition to the growing capsid. In this scenario, VP1 carrying motifs inserted in the PLA2 are not incorporated at all in the assembling capsid thus leaving more room for genome packaging, because the steric hindrance of internalized VP1u is removed. An alternative explanation for the increased packaging fitness is that the inserted motif forces VP1u to remain external, thus removing steric barrier to genome packaging. Our data provide support for the latter: pulldown with the respective affinity materials (e.g., GFP-agarose beads) showed significant enrichment if nanobody, mMobA, DCV, and SNAP tag were inserted into VP1u (residues 1:160), all of which improved also packaging fitness (Figure 4, Supplemental Figure 9). This means that these VP variants are incorporated in the capsid such the inserted motif is accessible on the capsid exterior. FLAG tag insertions into VP1u, which presumably did not interfere with internalization, did not increase pulldown fitness. Neither did WDV insertions, which is consistent with their deleterious impact on packaging fitness. Furthermore, we found that overall binding fitness was neutral (peaked around wildtype fitness, Figure 2 and Supplemental Figure 10), which was expected as most VP subunits (VP2 & VP3) contain the wildtype determinants of proteoglycan and AAVR binding (45, 65). However, we observed a notable drop for binding fitness for VP1u insertions of motifs that package well (e.g., nanobody; Pearson correlation coefficient −0.641), possibly due to steric hindrance of virus binding when it carries large external motifs. The only motifs that were not impaired for binding upon VP1u insertions are the FLAG tag, which can be explained by its small size, as well as SpyCatcher and WDV, which are motifs that our pulldown data suggest were not compatible with virus assembly when inserted into VP1u.

**Figure 3:**
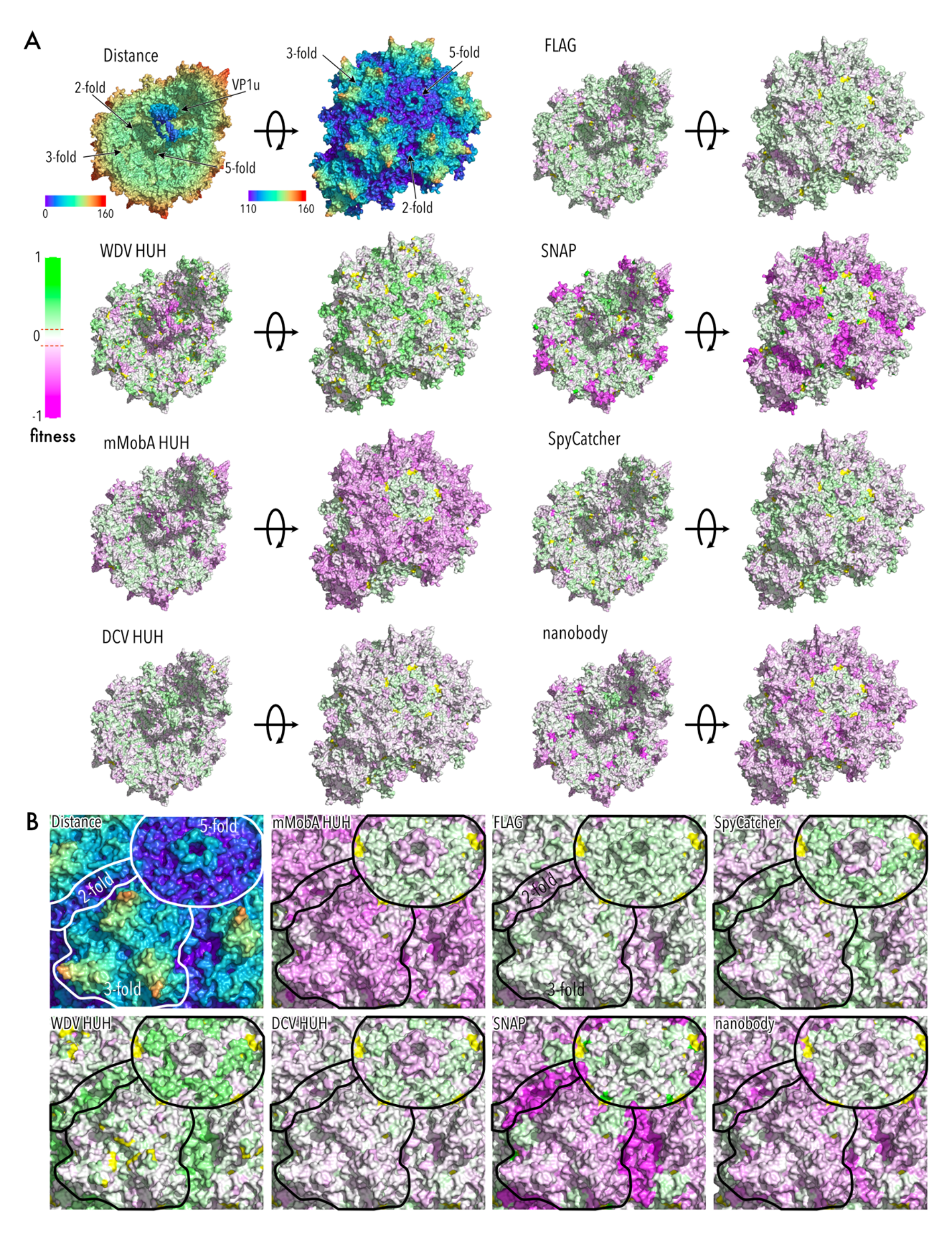
Packaging fitness of AAV domain insertion libraries mapped to the capsid structure. (A) Top left corner: AAV-DJ capsid structure view from the inside (left) and outside (right) radially color-cued. 2-, 3-, and 5-fold axes are indicated. VP1u domain was modeled using RoseTTAFold (98) and manually positioned. All other structures show packaging fitness heatmaps of the indicated domain insertions. Green indicates higher and magenta lower fitness than AAV-DJ (RCSB PDB 7KFR). (B) Zoom of the outside structures from (A). 2-, 3-, and 5-fold axes are outlined.

**Figure 4:**
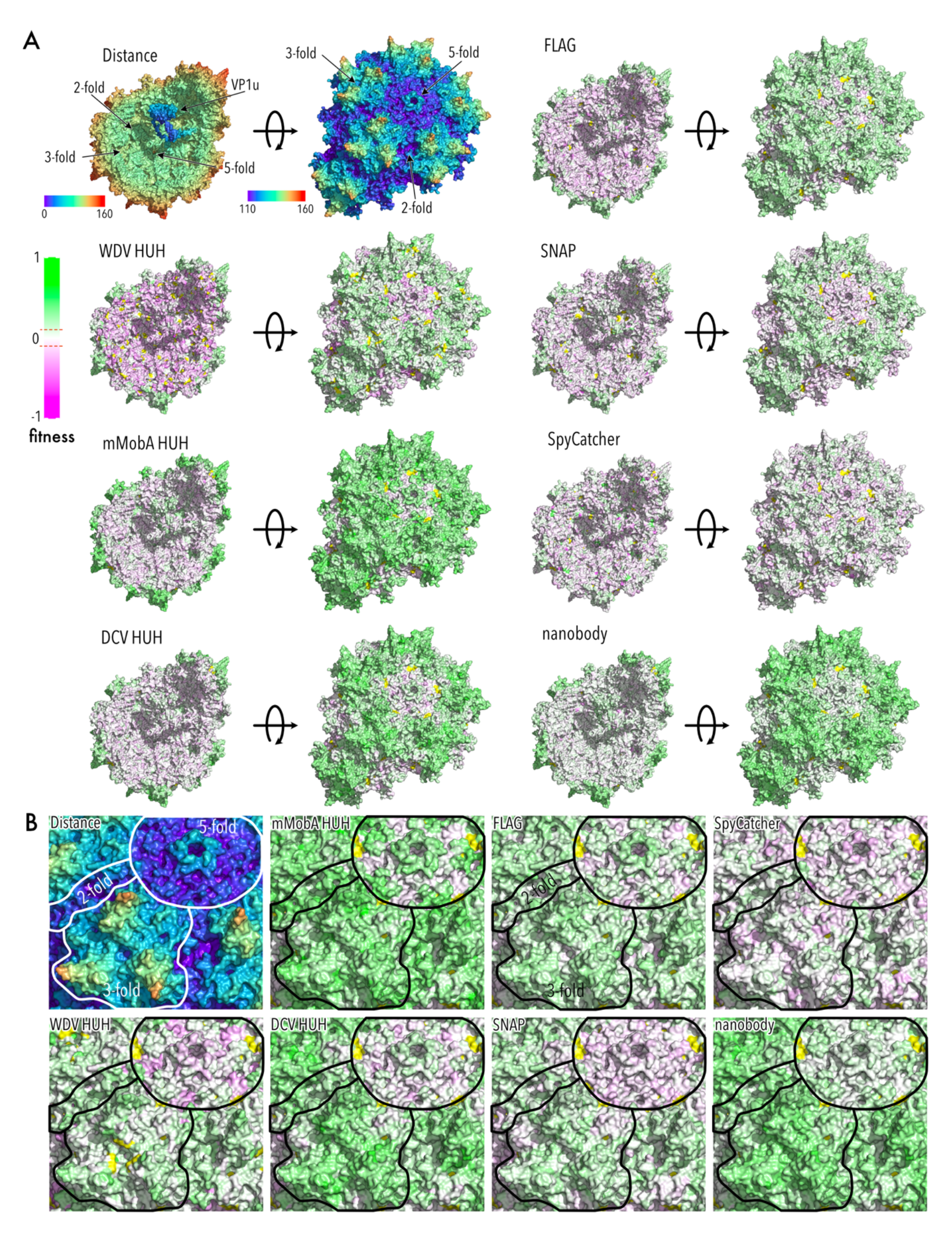
Pulldown fitness of AAV domain insertion libraries mapped to the capsid structure. (A) Top left corner: AAV-DJ capsid structure view from the inside (left) and outside (right) radially color-cued. 2-, 3-, and 5-fold axes are indicated. VP1u domain was modeled using RoseTTAFold (98) and manually positioned. All other structures show pulldown fitness heatmaps of the indicated domain insertions. Green indicates higher and magenta lower fitness than AAV-DJ (white) (RCSB PDB 7KFR). (B) Zoom of the outside structures from (A). 2-, 3-, and 5-fold axes are outlined.

Consistent with the role and required timing of VP1u in virus trafficking upon endocytosis, uptake fitness (assay measures presence of viral genome in any compartment inside the cell (66)) was impaired for all motifs inserted into the N-terminus of VP1 (Figure 2 and Supplemental Figure 11). It has previously been demonstrated that premature exposition of VP1u decreases infectivity (67), meaning that variants with pre-externalized VP1u escape less efficiently from the endosome and are degraded in the lysosomal compartment (23). We observed the same for infectivity fitness, with the notable exceptions of FLAG and WDV.

### Effects of domain insertions near AAV’s 3-fold axis

Insertion into protrusion near the 3-fold axis (residues 420-620; including VR4–8) generally impaired virus packaging (Figure 2 and Figure 3), which is consistent with the highly interdigitated structure of this region and its role in early assembly of VP trimers that are then forwarded to the nucleus as capsid building blocks (20, 36, 37). VP1 with insertions in this region may interfere with efficient trimer assembly (through a kinetic mechanism or by promoting off-pathway products), which would lower the overall capsid assembly efficiency. Nevertheless, several lines of evidence suggested that there appears to be some plasticity in trimer assembly to accommodate VP1 insertion variants so that they can be incorporated. For one, the overall hit to packaging fitness depended on the specific inserted motif. As expected, we saw that FLAG peptide insertions were relatively benign, but so were insertions of two HUH tags (DCV and WDV), and SpyCatcher (Figure 2 and Figure 3). There was no clear correlation with motif size hinting at more complex determinants for insertion fitness in this region. Second, despite impaired packaging fitness, pulldown fitness was greater than wildtype for all motifs, suggesting that they did become incorporated into purified AAV capsids, albeit at a lower overall efficiency (Figure 2 and Figure 4).

As for the N-terminus of VP1, binding fitness was not impaired for insertions in the 3-fold protrusion (Figure 2 and Supplemental Figure 10). In fact, the three HUH tags showed increased binding, which could be due to their generally higher cationic surface charge (Supplemental Figure 1B) aiding interaction with negatively charged components of the extracellular matrix, such as proteoglycans. Consistent with higher surface binding, we found higher uptake fitness than wildtype AAV-DJ in the case of DCV and WDV, when inserted into VR5 or VR8 (Figure 2 and Supplemental Figure 11). We note that binding and uptake fitness measured in this high throughput assay for insertion of mMobA into VR4 match our results from our previous engineering of this region (32). Despite higher binding and uptake, none of the insertions in this region could achieve wildtype infection efficiency (Figure 2), suggesting that domain insertion impacted later steps of virus trafficking to the nucleus.

### Effects of domain insertions near AAV’s 2-fold valleys

The neighborhood near the 2-fold symmetry center, which includes VR9, emerged as another region with distinct motif-specific phenotypes. This is consistent with earlier studies showing that dynamics of the 2-fold regions are essential for genome packaging (39) and AAV infectivity (68). Here, we observed both strongly deleterious fitness (e.g., SNAP tag) and strongly beneficial fitness (SpyCatcher and WDV). In all cases pulldown fitness was positive (Figure 4), suggesting the incorporation of at least one VP1 insertion variant that contains a motif insertion in this region. Interestingly, binding, uptake, and infectivity were impaired for most motifs.

### Unbiased clustering of insertion fitness reveals contiguous functional units in AAV capsids

Taken our fitness measurements across all motifs, all insertion positions, and all measured phenotypes in aggregate, we noticed patterns in fitness variance that appear correlated in contiguous regions of the AAV capsid. For example, packaging fitness varied predominantly in the PLA2 domain of VP1u and the 2-fold symmetry axis (Figure 5A). Focusing on residues unique to this interface, we found that packaging fitness distributions of interface and non-interface residues were not significantly different when DCV or FLAG peptide were inserted. However, fitness was significantly improved for mMobA, SpyCatcher, or WDV insertions into interface residues (Figure 5C). SNAP tag insertions were strongly deleterious. The variance at the 3-fold symmetry axis was markedly different. As described above, all motifs except WDV lowered packaging fitness, which resulted in lower overall fitness variance at this interface (Figure 5D). Remarkably, uptake fitness variance was greatest in the protrusion around the 3-fold axis and still considerably high along the 2-fold axis (Figure 5B). For all measured phenotypes, variance was relatively low around the 5-fold symmetry axis (Supplemental Figure 12).

**Figure 5:**
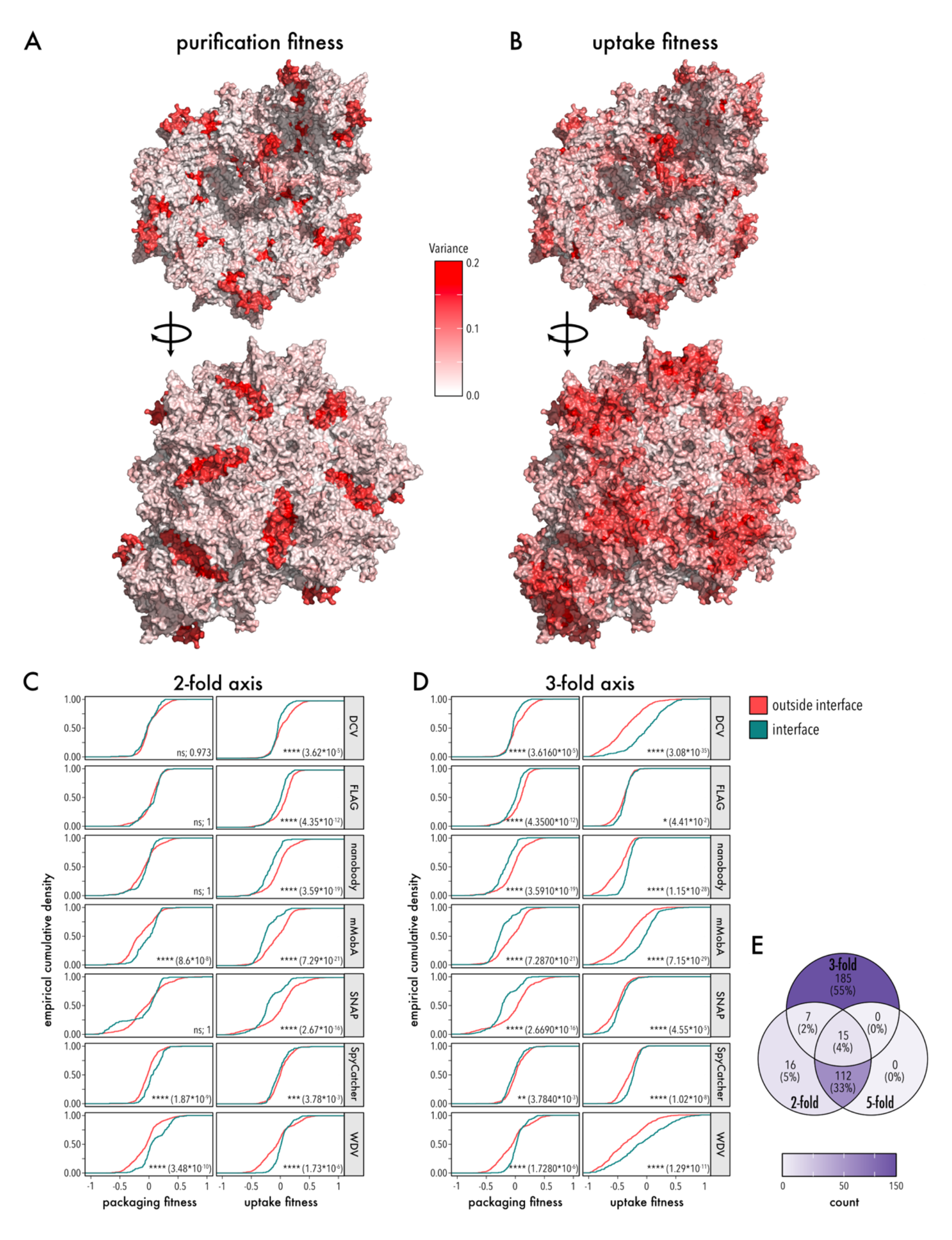
Variance of packaging and uptake fitness. (A,B) Variance of packaging fitness (A) and uptake fitness (B) from all domain insertion libraries mapped to the AAV capsid structure. The capsid inside (top) and the capsid outside (bottom) are shown (RCSB PDB 7KFR). (C) Empirical cumulative density insertional fitness of residues within (petrol green) and outside (red) the 2-fold axis (left) and 3-fold axis (right). Significance of distribution differences was tested using a two-sided, two-sample Kolmogorov-Smirnov test. Significance level and p values are shown. (D) Venn diagram showing which of the 335 AAV interface residues are unique and shared among interfaces.

If we think of different inserted motifs as different degrees of perturbation (e.g., weak for FLAG peptide insertion, strong for large nanobody insertion), then the positional and domain-type dependence of insertion permissibility we observed in our data suggests that we were measuring spatially resolved information of how different aspects of AAV fitness responds to these different degrees of perturbation.

To link insertion permissibility phenotypes to mechanistic structure/function relationships, we used an unbiased clustering approach (UMAP) (69). This resulted in five robust clusters that map to regions of the capsid with distinct roles in AAV biology (Figure 6A). Importantly, these clusters mapped to structurally contiguous (not interspersed) regions of the AAV capsid (Figure 6B,C).

**Figure 6:**
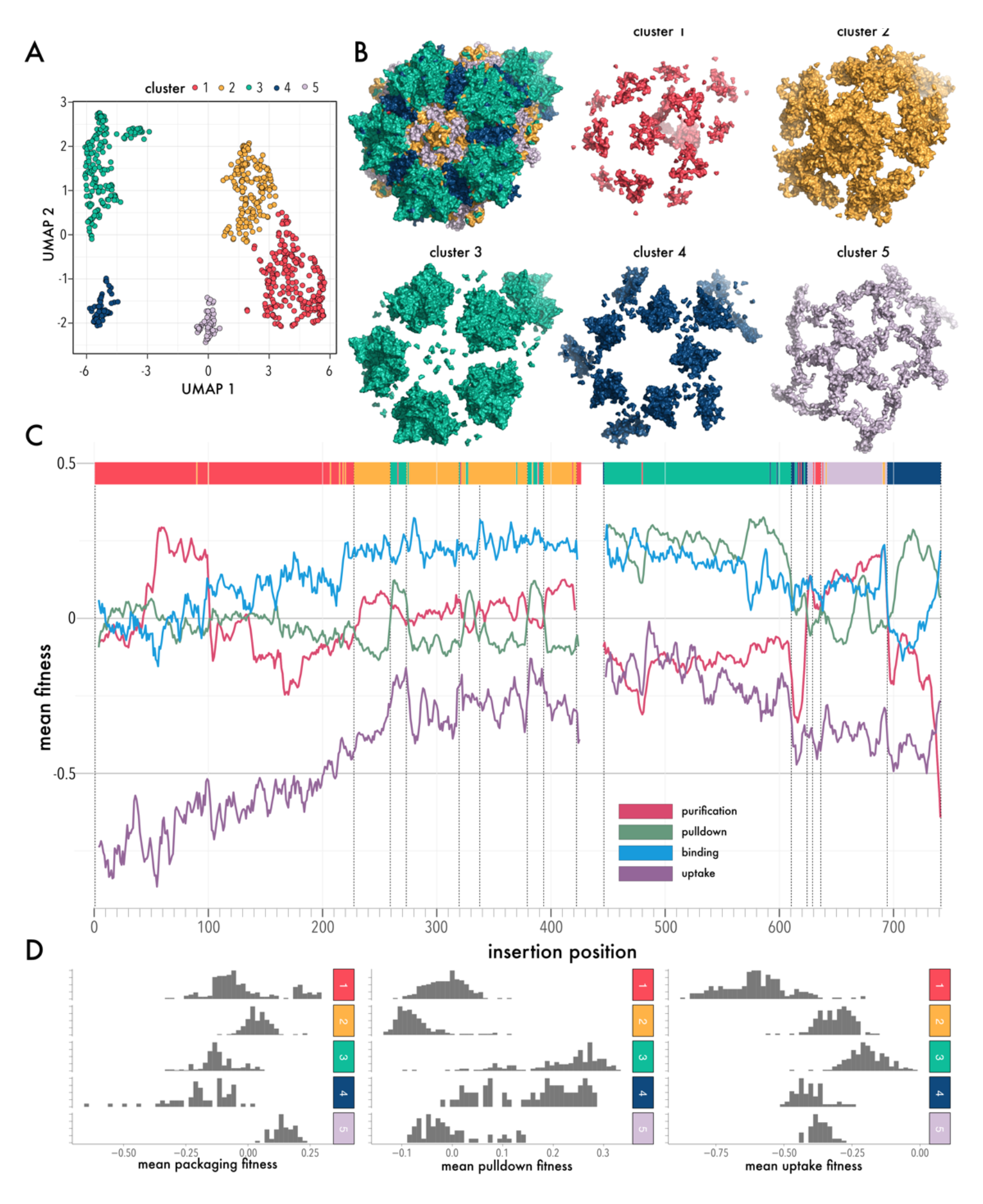
Unbiased clustering of insertion fitness. (A) UMAP cluster analysis of the AAV domain insertional profiling data resulting in five distinct clusters. (B) Cluster map to distinct capsid regions: (1) N-terminus of VP1 and bases of the 3-fold and 5-fold interface in red, (2) the HI-loop, the pore and the inner connecting residue layer in yellow, (3) protrusions of the 3-fold interface in green, (4) 2-fold axis in blue, and (5) the surrounding of the 5-fold pore in light lilac (C) Mean domain insertion fitness of packaging, pulldown, binding, and uptake shown by residues aligned to the UMAP clusters in (A). (D) Distribution of insertion fitness of packaging, pulldown, and uptake for each cluster.

Cluster 1 contains VP1u in addition to residues lining the bases of the 3-fold and 5-fold axes. Cluster 2 forms an extended network that comprises the HI loop and connects to the 3-fold axis protrusions. Cluster 3 represents protrusion at the 3-fold axis and residues on the external turns of the DE loop that line the pore at the 5-fold symmetry axis. Cluster 4 maps to the depression near the 2-fold symmetry axis and buried regions, which are part of the 3- fold axis. Cluster 5 predominantly maps to residues that line the capsid interior and the 5- fold pore, or that interdigitate HI loop, external residues of the 2-fold valley, and 3-fold protrusions (Figure 6B).

To understand the underlying mechanisms that drive clustering, we segregated fitness phenotype distributions by cluster identity (Figure 6D). Considering packaging fitness, we found that insertion into two clusters (1 and 5; containing VP1u and the network of residues that connect HI loop to the base of the 3-fold protrusion) were associated with improved packaging fitness, while clusters 3 and 4, which represent the interdigitated external region 3-fold-and 2-fold axes, were associated with poor packaging fitness. Insertions into capsid lining regions (cluster 2) were neutral with respect to packaging. With different measured phenotypes, these association patterns change: Considering pulldown fitness, we found that clusters representing buried residues or those lining the capsid interior (i.e., clusters 1, 2 & 5) were associated with poor fitness compared to those in externally accessible regions (cluster 3 & 4). For uptake, cluster 1 (which contains VP1u) had the worst fitness and cluster 3 (the 3-fold protrusion) was closest to wildtype fitness.

Using existing knowledge about structure and function of AAV, we can begin to make association between clusters and their specific roles in AAV packaging and infection. The strong association between poor pulldown fitness, uptake fitness, and cluster 1 is obvious considering it being comprised of mostly buried or internal residues and containing VP1u, which encodes the required PLA2 domain and nuclear localization signals (19, 21, 22, 46). Conversely, association with higher packaging fitness would be compatible with motif insertion promoting externalization of VP1u, thus decreasing steric hindrance with the packaged genome. The sensitivity of cluster 3 with respect to packaging efficiency is consistent with trimer formation and stability, which are key determinants of capsid assembly (36, 37). Most binding sites for cellular receptors (e.g., proteoglycan, AAVR) are located near the 3-fold axis, as well (14, 42–45, 65, 70). Flexibility of the 2-fold interface has previously been linked not only to AAV infectivity (68), but also genome packaging (39), which would explain how insertions may drive poor packaging fitness. The relatively neutral fitness of cluster 2 (e.g., packaging, uptake) is consistent with most insertions ending up on the capsid interior, not interfering with assembly or cellular uptake. While the 5-fold pore is part of this cluster, a numeric simulation of VP1 copy numbers ranging from 1-10 suggests that around half of the twelve 5-fold pores are assembled from non-VP1 only, and thus available as an alternative pathway for Rep-mediated genome packaging. Cluster 5 has an intricate structure, with the HI loops that surround the 5-fold pore like the blades of an aperture and connecting to the base of the 3-fold axis. Prior studies that have suggested a link between conformational changes at the 3-fold protrusion upon binding cell surface proteoglycan are communicated to conformational change at the 5-fold pore, priming the release of VP1u (71).

### Disulfide crosslinking to probe conformational flexibility

If the correlated conformational plasticity of clusters plays a role in packaging and/or infectivity, we would expect that changing conformational dynamics would impact these functions. One way to test this idea is by replacing two proximal residues by cysteines, such that a cystine disulfide link is formed once AAV is exposed to an oxidizing environment (i.e., after release from producer cells). If the two mutated residues are part of the same cluster of correlated conformational plasticity, and assuming that the individual cysteine substitution are benign, we expect a less significant effect on infectivity compared to if the two residues belong to different clusters. Our choices of residue pairs are summarized in Supplemental Figure 13A. Unlike domain insertions, which were only done in VP1, cysteine substitutions were introduced into all VPs. All single-and double-mutants produced near to wildtype titers (Supplemental Figure 13A; one-way ANOVA n.s.; Dunnett’s test with wildtype AAV-DJ as control: n.s.).

Some single cysteine mutants had dominant deleterious effects on infectivity (e.g., W608C, H292C). For double-mutants in the ‘within cluster’ set that had wild-type single mutant infectivity, infectivity was comparable to wildtype (e.g., H625C/Y426C, Supplemental Figure 13B), suggesting the minimal disruption of conformational dynamics. For pairs that belong to different clusters, all but one (H643C/Y350C) showed effects on infectivity that differed from what was predicted based on the individual single mutants (Supplemental Figure 13B). Several of these pairs involve interfaces that undergo conformational dynamics during infection.

For example, mutating F671C in the HI loop was strongly deleterious to infection, but this phenotype was rescued in the background of H255C, which by itself was benign. Prior studies have shown that heparin binding near the 3-fold and 2-fold axes induces an HI loop re-arrangement and an iris-like opening of the channel located at the 5-fold axis (71). HI loop deletions, substitutions, and insertions have shown that this loop, while flexible in amino acid composition and length, is critical for proper VP1 incorporation and infectivity (72). The same study showed that interaction of the HI loop with the underlying EF loop is mediated by hydrophobic pi-stacking interactions (F661/P373; in AAV2 numbering) and that disrupting this interaction lowers infectivity by preventing VP1 incorporating into assembled capsid. Our results are reminiscent of this mechanism. Positions F671/H255, which can form NH···π hydrogen bonds, are both conserved across AAV serotypes. H225C may be benign as this supports formation of an aromatic-thiol π hydrogen bond, thus allowing VP1 to incorporate or retaining structural rearrangement after receptor binding. Conversely, F671C is disruptive as it removes the aromatic component of π-stacking interactions, prevents VP1 incorporation and/or disrupts these rearrangements. The H255C/F671C double mutant may rescue infectivity by forming a disulfide bond to substitute as a stand in for π interactions.

In another example, H423C (at the base of the 3-fold axis) and V613C (with the 2-fold interface) individually had little effect, but together strongly impaired infectivity. This trend held true for adjacent pairs (H360C/437C; H428CL737C) that similar linked clusters comprising the 3-fold protrusion and 2-fold axis. Interestingly, several prior studies have linked AAV infectivity and conformational dynamics at the 3-fold and 2-fold axes. In addition to structural rearrangements in the HI loops, heparin binding to AAV2 causes significant rearrangement of 3-fold protrusions and the 2-fold valleys (71). Selective oxidation of tyrosine residue at the 2-fold dimer interface lowered infectivity (68). A mutation (R432A in AAV2) remodeled intramolecular and intermolecular hydrogen bond networks propagating from the 3-fold to both 2-fold and 5-fold axes (38, 40).

### Engineerable hotspots near the 2-fold axis and in the HI loop

We previously used the HUH tag mMobA in AAV-DJ VR4 to covalently link targeting scaffolds to the AAV capsid, which redirected AAV tropism *in vitro* (32). Here, we interrogated the entire AAV capsid for suitable HUH tag insertion hotspots. Comparing fitness maps for all three HUH tags, there were many differences amongst tags, which is likely related to their different biophysical properties (Supplemental Figure 1). We noticed that WDV was remarkably different two all other inserted domains in several regards. For one, packaging fitness was improved over wildtype when this domain was inserted into HI loops or along the 2-fold axis (Figure 3). Binding fitness was generally strong, but insertions into the 2-fold axis were deleterious (Supplemental Figure 10); this was the only insertion type for which we saw a deleterious phenotype for this assay. For uptake fitness, we observed a strong segregation in fitness between 3-fold protrusion and 2-fold and 5-fold axes (Supplemental Figure 11). Given that most previous studies have investigated VRs in the 3-fold protrusion for capsid engineering, we turned our attention to insertion sites near the 2-fold and 5-fold axes that had near wildtype packaging fitness in the NGS-based assay. We produced ten VP1 WDV insertion variants individually as crude cell lysates and measured titers, which all were comparable to wildtype (Figure 7A, one-way ANOVA n.s.). Testing each crudely enriched variant for the ability to infect HEK293FT cells, we found that several WDV variants inserted into surface exposed sites retained infection potency (Figure 7B). Among those were insertions into the 3-fold protrusions (S268), DE loop of the 5-fold pore (T331), HI loop (N664), and two sites along the 2-fold axis (Y702, K708). All sites had positive fitness in the NGS-based pulldown assay suggesting the WDV-VP1 does become incorporated into AAV capsid (Figure 2 and Figure 4). For two variants, N664 and K708 (Fig, 7C), we produced as iodixanol-gradient purified virus, which both trended to produce at higher titer compared to wildtype (Figure 7D) and incorporated VP1 as confirmed by Western blot (Supplemental Figure 14). As we have previously shown (32), HUH tags mediate the attachment of ssDNA antibodies to AAV, which in turn increased infectivity in cells that express, on the cell surface, the antigen recognized by the antibody. Using surface-expressed GFP (GFP-GPI) as a test case, we tested infectivity of the two purified WDV variants and wildtype AAV-DJ with and without conjugation to an ssDNA-linked anti-GFP antibody. Note that expression of GFP-GPI alone reduced cell health likely related to ER stress. Whereas infectivity of WDV variants was unaffected by conjugation to anti-GFP just like for wildtype AAV-DJ (expected as it was a mock conjugation since it does not contain a HUH tag), we saw a boost to infectivity for both N664-WDV and K708-WDV upon co-expression of surface expressed GFP (Figure 7E), suggesting that anti-GFP became conjugated to WDV and then enhanced infectivity by directing AAV toward surface expressed GFP as a binding receptor (two-way repeated measure ANOVA p-value 0.0063).

**Figure 7:**
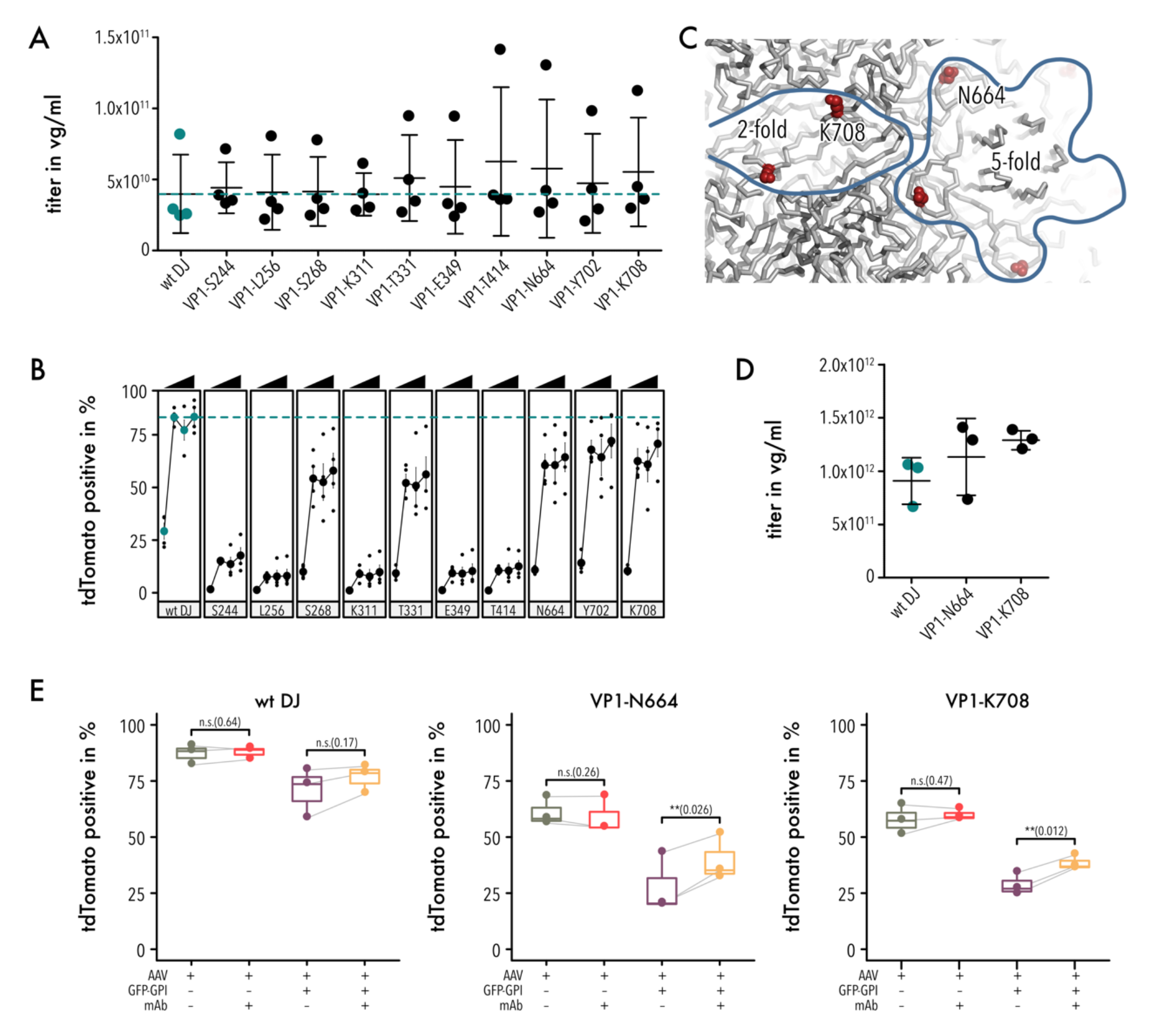
Fitness of WDV insertion variants. (A) Quantification of packaging titer of AAV-DJ and 10 different WDV insertion variants via qPCR. Data are means ± SD. (B) Infection fitness of the WDV insertion variants in (A) quantified by measuring the percentage of tdTomato positive cells 48 hours post transduction at the indicated MOIs. Data are mean ± SEM. (C) Zoom to the 2-fold and 5-fold axes (outlined) of the capsid surface. Positions of N664 and K708 are shown as red spheres. (D) Quantification of packaging titers of AAV-DJ and WDV insertion variants N664 and K708. Data are mean ± SD. (E) Infection fitness of the WDV insertion variants N644 and K708 quantified by measuring the percentage of tdTomato positive cells 48 hours post transduction at an MOI of 1×10^4^ vg. If indicated, cells were co-expressing GFP-GPI and/or a ssDNA-anti-GFP antibody was added. Data shown as box plots. Lower and upper hinges of boxes indicate 25^th^ and 75^th^ percentile, respectively. Mean is indicated by a horizontal bar in each box. Whiskers extend 1.5 * IQR. Two-way repeated measure ANOVA was used to test the significance of variance between means (AAV-DJ, N664, K708) for conditions with and without GFP present. Presence of the mAb was a significant source of variation when GFP-GPI was expressed (p value 0.0063) but not without (p value 0.514). Significance levels and p values for pairwise comparisons using a Bonferroni correction are shown.

## DISCUSSION

Viral vectors are an essential component for gene delivery in therapies treating inherited disorders and cancer. Adeno-associated virus (AAV), in particular, is widely used in both approved therapies and in ongoing clinical trials because of its good safety in humans and ability to drive long-term expression in both dividing and non-dividing cells. Unfortunately, several of the evolved properties of AAV are mismatched to clinical needs (e.g., broad tropism, limited payload capacity, existing serum-immunity in most of the human population) or they pose biomanufacturing challenges (e.g., scale up of helper virus free production, yield of full virions to maximize potency).

Motivated by these challenges, there have been extensive efforts to re-engineer AAV properties in the past including directed evolution approaches, such as repeated mutagenesis, capsid shuffling (73), viral display of short peptides (52–54), and adding larger, structured targeting scaffolds, such as antibodies, nanobodies, DARPins, or affibodies (30–33, 74). Most of these studies have focused on regions in the 3-fold protrusion, commonly VR8, VR4, or VP termini. Recently, deep mutagenesis of the entire capsid protein, combined with machine learning (8, 47, 75), demonstrated that learned sequence & function relationships can aid the prediction of sequence variation to improve desired AAV traits. However, deep mutagenesis in AAV has so far been limited to amino acid substitutions.

Here, we combined the concept of deep mutagenesis with scaffold insertion to systematically measure the fitness of AAV containing VP1 with seven different motifs inserted in between every two residues. This comprehensive analysis quantitatively links where different structured motifs can be inserted into VP1 to retain compatibility with AAV virion assembly and packaging, cell binding, and uptake. The fitness profiles show that there is a strong dependence on insertion position and type of inserted motif, with several regions showing diverging fitness for different measured phenotypes (e.g., VP1u for packaging vs. cell uptake, Figure 2). This highlights that the outcome of sequence variation can have multi-facetted impact on AAV properties, and it calls for integration of assays across several clinically relevant AAV attributes to safeguard against inadvertent optimization for undesired traits. In recent years, we have gained broad access to precision variant library engineering (35, 76, 77), Next-Gen sequencing (enabling counting number of sequence variants in a highly diverse population before and after applying a test for fitness (62)), unified analytical frameworks to interpret these large dataset (78), and machine learning approaches for deciphering sequence/function relationships (79). Taken together, generating and interrogating large AAV variant libraries has become feasible. Our focus can now shift towards what these datasets tell us about AAV biology and how they can guide and accelerate viral vector engineering.

For example, while fitness of many insertion sites is consistent with known AAV structure and function (e.g., the importance of trimer assembly along 3-fold axis in capsid assembly (36)), the high packaging fitness when motifs are inserted into the PLA2 domain of VP1u was surprising. Several studies have elucidated VP1u and VP2 dynamics as part of events after virus uptake by the cell, in which the internalization of VP1u PLA2 is a required step for endosomal escape (21, 46). VP1u internalization during maturation of AAV virions is less well understood but appears to be coordinated with genome packaging (39). The existence of such mechanism would reconcile recently proposed models of stochastic assembly (13) (which should result in both internalized and externalized VP1u) and the well known requirement for heating or pH lowering to induce VP1u externalization before it can be detected by VP1-specific antibodies (17, 19, 72, 80). It is plausible that domain insertions interfere with this process, leaving VP1u externalized, which would leave more room inside the AAV capsid for genome packaging by Rep78 due to lack of steric hindrance (one VP1 occupies 85×10^3^ Å^3^ or 1/35^th^ of the available space inside the capsid) but at the loss of infectivity. Interestingly, VP1u and VP2 of the related parvovirus B19V have acquired an receptor-binding domain insertion just upstream of PLA2 and are always external (81–84), supporting the idea that addition of extra domains into VP1u leave it externalized.

Given that precise timing of VP1u externalization in the correct endosomal compartment is required for maximum infectivity (premature externalization reduced infectivity (67)), this may point to an evolutionary mechanism that balances virion packaging efficiency with infectivity. If the full/empty ratio is fundamentally constrained by a packaging/infectivity balance, this would have implications for biomanufacturing of AAV in which one of the major ongoing efforts is to find ways to enrich full particles. One prediction of our hypothesis is that full particles may contain fewer VP1, on average, compared to empty particles, and this is negatively impacting infection potency. The observation that overexpression of VP1 inhibits rAAV packaging is consistent with this idea (85). There may exist a Goldilocks regimen, just the right copy number of VP1, that maximizes both - genome content and endosomal escape. Further research is required to fully test this hypothesis, including measuring VP stoichiometry and genome content at the single capsid level.

Unbiased clustering of insertion fitness across several phenotypes also revealed a topological organization of AAV into regions that can be linked to correlates in AAV capsid assembly, genome packaging, and infectivity. Given that many of these roles have a basis in distinct regimes of capsid stability and dynamics, we hypothesize that inserting different domains (which represent different degrees of perturbation) probes conformational plasticity at or near the insertion site. Put simply, it probes whether the insertion site is conformationally rigid (allowing no insertions) or conformationally flexible (allowing some or all insertions). Clusters emerge because conformational plasticity of different capsid regions impinges differently with different measured phenotypes (e.g., capsid flexing required for efficient genome packaging (39) vs. flexing during externalization of VP1u/PLA2 after cell uptake, which only happens in full, but not empty capsids (17)).

Similar ideas of spatially contiguous protein regions linked to specific functions have been proposed in the past, including protein “sectors” mapped through measuring amino acid co-evolution (86, 87), regional conformational flexibility mapped by circular permutation profiling (88) and domain insertion (49, 50, 76), or revealing the functional architecture of an enzyme from high-throughput enzyme variant kinetics (89). At their core, all these approaches use mutations to perturb sequence/function relationships. Similarly, by perturbing VP1 through domain insertion and measuring how it responds (in terms of assembly, infectivity, etc.), we learn how AAV structure intersects with AAV function. Going forward, domain insertional profiling in the background of different genetic backgrounds (i.e., serotypes) may further separate general principles of AAV assembly, function and serotype-specific properties (e.g., stability, immune evasion, tropism).

Our systematic domain insertion approach also revealed new opportunities for viral engineering. Following the intuition that engineering non-conserved, surface exposed, and tropism-determining loops is the likeliest path to change AAV properties, much of AAV engineering so far has focused on N-termini of capsid proteins, or variable loops of the 3- fold protrusions (30, 32, 34, 74, 90–94). However, systematic studies we and others conducted (50, 88), suggest that this intuition may be misleading. We found that permissibility to domain insertion is not correlated with conservation, surface exposure, or other static, structural features. Instead, dynamic features, such as regional flexibility, are predictive as to where a domain insertion is tolerated. In this study, we identified two new regions near the 2-fold and 5-fold axes that can tolerate the insertion of HUH tags (15kDa), which in turn enable the covalent linkage of antibodies (150kDa). In a proof-of-principle, we found that insertions have little impact on production levels and provide a modest boost to infecting cells that express the antibody’s cognate antigen. Further research and engineering are required to fully leverage the potential of these new engineerable hotspots. They represent an exciting opportunity to sidestep the constraint of directed evolution of targeting the region near the 3-fold axis. As this region is important for receptor binding, it is also the most antigenic region. In fact, the binding sites for several proteoglycans, AAVR, and neutralizing antibodies (42, 43, 45, 70 overlap. Approaches that shuffle the sequence of this region must apply selection pressure to selectively remove antibody binding whilst retaining the mode of cell binding and uptake. By providing an alternative site to which targeting scaffold can be linked, it may be possible to address this challenge more effectively.

## MATERIALS AND METHODS

### Cloning and library generation

All plasmids and libraries used in this study are listed in Supplemental Table 1 and were generated either by classical restriction enzyme cloning or Golden Gate Assembly (95). Restriction enzymes were obtained from NEB, standard oligos, labeled oligos as well as gBlocks from IDT, and oligo pools for library cloning from Agilent Technologies. For amplification of DNA sequences for cloning and NGS the PrimeSTAR Max DNA polymerase (Takara Bio Inc.) and for colony PCRs the OneTaq Quick-Load Master Mix Polymerase was used (NEB). PCR products were analyzed on 1 % TAE agarose gels, cut out and purified using the Zymoclean Gel DNA Extraction Kit (Zymo Research) by following the manufacturer’s instructions. Post cloning, plasmids were transformed into NEB^®^ Stable Competent *E. coli* cells and libraries into MegaX DH10B T1R Electrocomp Cells (Thermo Fisher), before plated on LB plates containing either carbenicillin (100 µg/ml) alone or a combination with chloramphenicol (25 µg/ml) depending on the selection marker(s) on the plasmids and libraries. To assess coverage of libraries a small amount from the transformed cells was taken, serial dilutions prepared, and plated on LB plates with the respective selection marker(s). The next day, colonies were counted to estimate coverage and colony PCRs were run to verify library diversity. Plasmids were isolated using the Zyppy Plasmid Miniprep Kit, ZymoPURE II Midiprep Kit or the ZymoPURE II Maxiprep Kit (all Zymo Research).

For the generation of AAV-DJ insertion libraries an altered cap-DJ gene sequence was used with mutated start sites for VP2 (T138A) and VP3 (M203K, M211L, and M235L), a T176A mutation eliminating the BsmBI cutting site, and a replacement of the HBD domain (R587- R590) by an HA-tag (AYPYDVPDYAA). The libraries were created using SPINE as previously published by us (35). In brief, the *cap* DJ-VP1 sequence was split up into 14 fragments, and oligos containing a genetic handle behind every amino acid position were designed. Every oligo contained barcodes for amplification, matching BsmBI restriction sites to assemble the 14 fragments, and BsaI restriction sites to swap out the handle. The genetic handle was first replaced by a chloramphenicol expression cassette flanked by BsmBI cutting sites to insert a selection marker for library presence. Next, the library was transferred into an AAV plasmid backbone encoding an EFS-driven miRFP670nano sequence terminated by a SV40-polyA, a p40 promoter with BsmBI sites for the library insertion and flanked by ITRs. Last, the chloramphenicol was replaced by different domains (nanobody, SpyCatcher, SNAP, mMobA, WDV or DCV) or a FLAG-tag. While the domains had 5aa SGGGG-domain-GGGGS linkers, the FLAG-tag was flanked by short SG-FLAG-GS linkers only. The DJ-VP1 silent mutation plasmid, that was used as a reference, was designed by introducing ten silent mutations into the VP1-only DNA sequence of AAV-DJ. To this end, codons for either arginine, serine or leucine were altered by two nucleotides each at positions that were 200- 300 bp apart from each other. The silent mutation VP1 coding sequence was ordered as a gBlock and cloned into the same backbone as the plasmid libraries, i.e., miRFP670nano expression cassette and a p40 promoter to drive VP1 expression and flanked by ITRs. Cysteine point mutations were introduced by site-directed mutagenesis of the wildtype rep2- capDJ plasmid. WDV insertion variants were generated by inserting the domain into the above mentioned p40-driven and altered AAV-DJ-VP1-only sequence by Golden Gate Cloning.

### Tissue culture

HEK293FT cells (Invitrogen) and 293AAV cells (Cell Biolabs) were maintained in DMEM (Gibco) containing 4.5 g/L D-glucose, L-glutamine, 110 mg/L sodium pyruvate, and supplemented with 10 % fetal bovine serum (Gibco) and 100 U per mL penicillin/100 μg per mL streptomycin (Gibco). Cells were kept in a humidified cell culture incubator at 5 % CO_2_ and 37 °C and passaged every 2-3 days when reaching 70-90 % confluency. For experiments with HEK293FT cells, plates were pre-coated with growth factor reduced basement membrane matrix Matrigel (Corning) prior to seeding. If applicable, HEK293FT cells were transfected with a plasmid encoding GFP-GPI using Turbofect (Invitrogen) while seeding and according to the manufacturer’s protocol. The amounts of DNA used for transfection are further specified in the sections of the different assays. For AAV productions, 293AAV cells were used only till reaching passage 10.

### AAV crude lysate production

293AAV cells were seeded into 6-well plates at a density of 500,000 cells per well. The next day, cells were transfected with 2.5 µg DNA using PEI and an equimolar ratio of the plasmids necessary for the respective AAV production. Three days post transfection, cells were harvested by flushing off the cells by pipetting. Cells were washed with PBS once and then subjected to five freeze and thaw cycles by alternating between liquid nitrogen and a 37 °C water bath. Cell debris was pelleted by centrifugation at 18,000 rpm at 4 °C for 10 minutes and the supernatant containing the AAV particles, was stored at −20 °C until use.

### Purified AAV production

Large scale productions of AAV were either done by the University of Minnesota Viral Vector and Cloning Core using a sucrose gradient or by ourselves following published iodixanol gradient density protocols (41, 96). In brief, four million 293AAV cells were seeded into 15 cm dishes and transfected using PEI and 47 µg total DNA per dish 48 hours post seeding. For the AAV-DJ library control, an equimolar triple-transfection was used composed of an Adeno-helper plasmid, a plasmid encoding the *rep2* and *capDJ* genes, and a transgene plasmid encoding an EFS promoter-driven miRFP670nano sequence flanked by ITRs. For AAV-DJ insertion library productions, a plasmid ratio of 1:0.1:0.1 of an Adeno-helper plasmid, a plasmid encoding rep2 and only VP1 of capDJ, as well as the respective AAV-DJ insertion library in which the AAV-DJ silent mutation variant was spiked in was transfected. To top up to 47 µg total DNA, a pUC19 stuffer plasmid was added. 72 hours post transfection, cells were detached with a cell lifter and cells pelleted by centrifugation at 400 xg for 15 minutes. Cell pellet was washed once with PBS and resuspended in a buffer containing 2 mM MgCl2, 0.15 M NaCl and 50 mM Tris-HCl at pH 8.5. Cells were cracked open using five freeze and thaw cycles. Free genomic and plasmid DNA was digested with a Benzonase Nuclease (Sigma-Aldrich). Cell debris was removed by centrifugation and, subsequently, the lysate transferred into ultracentrifugation tubes (Beckman Coulter). The iodixanol discontinuous gradient (15%, 25%, 40%, and 60% iodixanol concentration) was layered underneath the cell lysate. Density gradient centrifugation was done at 50,000 rpm for two hours at 4 °C using a 70.1 Ti rotor (Beckman Coulter). Post centrifugation, the 40% iodixanol phase containing the AAV was isolated, aliquoted, and stored at −80 °C until use. For pulldown, binding and uptake, as well as the DSF assays, AAV samples were dialyzed to PBS supplemented with 5% glycerol using 10 kDa Amicon Ultra-15 Centrifugal Filter Units (MilliporeSigma).

### qPCR

To determine the titer of crude lysate AAV samples, 1-5 µl of the crude lysate were mixed with PBS supplemented with 2 mM MgCl_2_ to a final volume of 50 µl. Then, 0.1 µl ultrapure Benzonase Nuclease (Sigma-Aldrich) was added. Samples were incubated for 30 minutes at 37 °C to digest non-encapsidated DNA. Next, 5 µl of a 10x Proteinase K buffer (100 nM Tris-HCl, pH 8.0, 10 mM EDTA, and 10% SDS) and 1 µl Proteinase K (20 mg/ml; Zymo Research) were added to stop the DNA digest and start the protein digest to free the ssDNA from the AAV particles. Samples were incubated for 20 minutes at 50 °C, followed by a heat inactivation of the enzymes for 5 minutes at 95 °C. The DNA was purified using the DNA Clean & Concentrator-5 Kit (Zymo Research) according to the manufacturer’s instructions for ssDNA purification. For purified AAV samples, the Benzonase digest step was skipped and only the Proteinase K and DNA purification steps were performed. All samples were diluted 1:1,000 in H_2_O prior to qPCR. The viral genome (vg) quantification was done on a QuantStudio5 Real-Time PCR System (Applied Biosystems) using the PowerUp SYBR Green Master Mix (Applied Biosystems) and following the manufacturer’s instructions. To calculate the viral titer (vg/ml) a plasmid standard with a known concentration of plasmid copies was used. Primer sets binding either the CMV-enhancer or the p40 promoter of the AAV genomes, as well as in the plasmid standard were selected (Supplemental Table 3).

### Pulldown assay of libraries

Between 1e9 and 1e10 vg were used as input material to bind to different magnetic beads for pulldown assays: SNAP-Capture Magnetic Beads (NEB) for SNAP tag insertion; Pierce Anti-DYKDDDDK Magnetic Agarose (Thermo Scientific) for FLAG tag insertion; Streptavidin Magnetic Beads (NEB) for nanobody, SpyCatcher, and HUH tag insertions. 80 µl bead slurry for SNAP pulldowns and 50 µl bead slurry for FLAG pulldowns were washed three times with 300 µl wash buffer I (0.15 M NaCl, 20 mM Tris-HCl pH 7.5, 1 mM EDTA). Then AAV libraries were mixed with wash buffer I and the beads to a final volume of 300 µl, before incubated on a slow shaker for 30 minutes at room temperature. Beads were washed again three times with 300 µl wash buffer I for SNAP beads and with PBS (pH 7.4, Gibco) for FLAG beads to remove unbound AAV particles, and finally resuspended in 50 µl PBS. For pulldown assays with streptavidin beads, 100 µl bead slurry was washed three times with 300 µl wash buffer II (0.15 M NaCl, 20 mM Tris-HCl pH 7.5, 1 mM EDTA). For nanobody and SpyCatcher binding, beads were pre-incubated with either 320 pmol biotinylated superfolder-GFP or 1,000 pmol biotinylated SpyTag, respectively, for 30 minutes on a slow shaker at room temperature in a total volume of 300 µl in wash buffer II. Afterwards, unbound superfolder-GFP and SpyTag were removed by washing the beads three times with 300 µl wash buffer II. For HUH tag pulldowns, AAV libraries were first reacted with 1 nmol biotinylated ssDNA oligos (sequences are given in Supplemental Table 3) in PBS supplemented with 0.05 % v/v salmon sperm DNA (Invitrogen), 1 mM MgCl_2_, and 1 mM MnCl_2_ for 15 minutes at 37 °C. Next, beads were incubated with AAV libraries for 30 minutes on a slow shaker at room temperature, before unbound AAV particles were removed by washing three times with 300 µl wash buffer II. Beads with bound AAV particles were resuspended in 50 µl PBS. Viral genomes of samples after the pulldown were purified using the Quick-DNA Microprep Plus Kit (Zymo research) and by following the manufacturer’s protocol.

### Binding and uptake assays of libraries

The binding and uptake assays were performed as previously described in Berry et al. 2017 (66), but using HEK293FT cells. In brief, 375,000 HEK293FT cells were seeded into 6-well plates using 2 ml culturing media per well. The next day, cells were incubated for 30 minutes at 4 °C. Afterwards, the media was aspirated and 200 µl cold DMEM containing AAV particles at an MOI of 1e4 added. The cells were further incubated for one hour at 4 °C. Next, cells were washed three times with ice-cold PBS to eliminate unbound AAV particles. For the binding assay, 150 µl ice-cold PBS was added, the cells detached with a cell scraper, and the cell suspension transferred to a microcentrifuge tube. For the uptake assay, 1 ml of pre-warmed culturing media was added immediately after the PBS wash and the cells incubated for two hours in a cell culture incubator to allow for uptake of the AAV particles. Then, the cells were detached by trypsinization and collected in a microcentrifuge tube, before washed three times with 200 µl PBS. The DNA from binding and uptake samples was purified using the Quick-DNA Microprep Plus Kit (Zymo research) and by following the manufacturer’s protocol.

### Infectivity assay of libraries

150,000 HEK293FT cells were seeded into 12-well plates using 1 ml culturing media per well. The next day, cells were transduced with purified AAV (in PBS supplemented with 5 % glycerol) at an MOI of 2e5 vg/cell. To this end, the media was aspirated, cells were washed with 500 µl PBS once. Purified AAV were mixed with DMEM without supplements to a final volume of 500 µl and added onto the cells. After two hours of incubation in the cell culture incubator, 1.5 ml culturing media (with supplements) was added. 24 hours post transduction, the temperature was reduced to 33 °C to promote protein expression rather than cell growth (97). 72 hours post transduction cells were prepared for cell sorting as follows. Media was aspirated, cells were washed with 500 µl PBS, 500 µl Accutase solution (Sigma-Aldrich) was added and incubated at room temperature until all cells detached. Cell suspension was transferred to a microcentrifuge tube and centrifuged for three minutes at 400 xg to pellet cells. Cells were washed two times with 500 µl PBS, before resuspended in 650 µl cell sorting buffer (PBS supplemented with 5 mM EDTA and 2.5 % FBS) and passed through a 35 µm cell strainer to avoid cell clumps. Cell sorting was performed by the University of Minnesota Flow Cytometry Resource (UFCR) on a FACS Aria II instrument (BD Biosciences) with a 85 µm nozzle by sorting miRFP670nano positive (excitation 640 nm laser, emission 670 nm/ 30nm bandpass filter). Post sorting, the DNA was extracted from the cells using the Quick-DNA Microprep Plus Kit (Zymo research) and by following the manufacturer’s protocol.

### NGS preparation

Purified DNA samples (library plasmid DNA, ssDNA from purified AAV, and DNA extracted after pulldown, binding, uptake, and infectivity assays) were amplified using primers binding 50 base pairs up and down stream of the VP1 coding sequence (Supplemental Table 3). For amplification, the PrimeSTAR Max DNA Polymerase (Takara Bio Inc.) was used according to the manufacturer’s recommendation with a 25 µl reaction volume, an annealing temperature of 62 °C and an elongation time of 15 seconds. At least five reactions were pooled for each DNA sample, whereas the cycle number was kept at a minimum to obtain >50 ng per sample. PCR products were purified using the DNA Clean & Concentrator-5 kit (Zymo Research) and by following the manufacturer’s instructions for dsDNA purification. The DNA was eluted in a 10 mM TRIS buffer with pH 8.0. Prior to sequencing, the DNA was quantified using the Qubit 1X dsDNA HS assay kit and a Qubit 4 Fluorometer (both Invitrogen), and the DNA of the insertion libraries from the same assay was pooled in an equimolar ratio. >50 ng of each DNA pool was submitted to the University of Minnesota Genomics Center, where Nextera XT libraries were created, and samples sequenced using a NovaSeq SPrime 150 paired end run.

### Sequencing data analysis and enrichment calculation

Forward and reverse reads were aligned individually using a DIP-seq pipeline (76), slightly modified for SPINE compatibility and for updated python packages. The code for handling data from domain insertion library sequencing is available at: https://github.com/SavageLab/dipseq. This pipeline results in .csv spreadsheets (available as processed data, along scripts to reproduce manuscript figures, which are available at https://github/com/Schmidt-lab/AAV_Insertion_Profiling) indicating insertion position, direction, and whether it was in frame. Fitness was calculated from the frequency of a given VP1 variant (*i*) after packaging (*s*) relative to the frequency of that variant in the input library (*u*), normalized to wildtype AAV (*wt*):

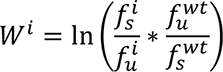

Fitness standard error for each variant was calculated assuming a Poisson distribution.

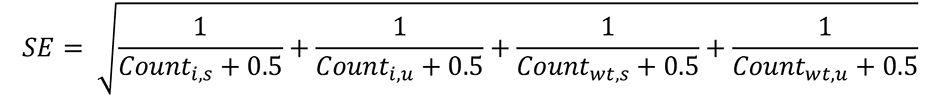

### Western blot

For VP protein analysis 3×10^9^-1×10^10^ vg of purified AAV were mixed with 12.5 µl 4x Laemmli Sample Buffer (Bio-Rad, supplemented with 10% 2-mercaptoethanol) and topped up to a final volume of 50 µl with PBS. Samples were denatured for 10 minutes at 95 °C and afterwards chilled on ice. Protein samples, and 5 µl of the Precision Plus Protein Dual Color Standard (Bio-Rad), were separated by molecular weight on a 7.5 % precast polyacrylamide gel (Bio-Rad) in Tris/Glycine/SDS Electrophoresis Buffer (Bio-Rad) for 85 minutes at 120 V. Next, proteins were transferred to a nitrocellulose membrane (pore size 0.45 µm; Thermo Scientific) in an ice-cold blotting buffer (25 mM Tris Base, 96 mM glycine, 20 % methanol) for 80 minutes at 110 V. The membrane was washed once in TBS-T (20 mM Tris Base, 137 mM NaCl, pH 7.6, 0.05% Tween-20) and incubated in 5 % skim milk solution in TBS-T for two hours at room temperature on a slow shaker to block non-specific binding. A primary antibody detecting either all three VP proteins (anti-AAV VP1/VP2/VP3 mouse monoclonal, B1, supernatant, Progen) or only VP1 (anti-AAV VP1 mouse monoclonal, A, lyophilized, purified, Progen) was diluted 1:250 in 5 % skim milk solution in TBS-T and incubated overnight at 4 °C. The next day, the membrane was washed four times for 5 minutes in TBS-T on a shaker, before the secondary antibody was added (anti-mouse IgG-peroxidase antibody produced in goat, Sigma-Aldrich). The secondary antibody was diluted 1:50,000 in 5% skim milk solution in TBS-T and incubated for two hours at room temperature on a slow shaker. Afterwards, the membrane was washed again four times for 5 minutes in TBS-T at room temperature to remove unbound antibodies, before the SuperSignal West Dura Extended Duration Substrate kit solution (Thermo Scientific) was applied and incubated for two minutes at room temperature. The chemiluminescence signal was detected with an Amersham Imager 600 (GE Healthcare) using exposure times between one second and ten minutes, depending on the signal intensities. Quantification of VP expression was done using ImageJ.

### Differential Scanning Fluorimetry (DSF)

First, 5,000X SYPRO Orange dye (Invitrogen) was diluted 1:100 in PBS with 5 % glycerol. Then, each sample was prepared by mixing 50X SYPRO Orange dye 1:10 with >5×10^9^ vg of purified AAV samples in PBS with 5 % glycerol to a final volume of 25 µl or 50 µl. Samples were mixed by pipetting up and down and pipetted into a 0.1 ml MicroAmp Fast Optical 96- Well Reaction Plate (Applied Biosystems). The plate was sealed with an optical adhesive cover (Applied Biosystems) and spun down for two minutes at 1,000 xg. A melt curve experiment was run on a QuantStudio5 Real-Time PCR instrument (Applied Biosystems) using the x1-m4 filter set (excitation filter: 470 nm/ 15nm, emission filter: 623 nm/ 14nm) and the following settings: 30 °C for two minutes, temperature increase from 30 °C to 99 °C in 0.5 °C and 30 seconds increments, and a final incubation step of two minutes at 99 °C. Lysozyme at a concentration of 0.1 mg/ml with a determined melting temperature of 70 °C was used as a reference control within each run. Post processing of the melt-curve data was done in MATLAB R2021a (The Mathworks Inc., Natick, Massachusetts). A smoothing spline (smoothing parameter p=0.9) was fitted to the data before calculating the numerical gradient ∂(Fluorescence)/∂(Temperature).

### Transmission electron microscopy

To quantify the empty to full capsid ratio, negative staining and transmission electron microscopy was performed by the Characterization Facility, University of Minnesota. At least 300 AAV particles were manually counted per sample.

### Infectivity assay of cysteine mutants and WDV variants

75,000 HEK293FT cells were seeded per well of a 24-well plate using 0.5 ml culturing media. If applicable, cells were transfected with 100 ng GFP-GPI plasmid while seeding. The next day, media was exchanged, and cells transduced with crude lysates or purified AAV at the indicated MOIs. 48 hours post transduction cells were prepared for flow cytometry as follows. Media was aspirated, cells were washed with 500 µl PBS, 250 µl Accutase solution (Sigma-Aldrich) was added and incubated at room temperature until all cells detached. Cell suspension was transferred into a microcentrifuge tube and centrifuged for three minutes at 400 xg to pellet cells. Cells were washed two times with 300 µl PBS, before resuspended in 600 µl flow cytometry buffer (PBS supplemented with 5 mM EDTA and 2.5 % FBS) and passed through a 35 µm cell strainer to avoid cell clumps. Flow cytometry was performed either on a LSRFortessa X-20 or a FACSymphony A3 Cell Analyzer (both BD Biosciences) equipped with 561 nm and 488 nm lasers to detect tdTomato and GFP positive cells, respectively. Minimum 10,000 single cell events were recorded per sample. Data analysis was performed using the FlowJo 10.8.0 software (BD Biosciences).

## DATA AVAILABILITY

Plasmids and plasmid libraries are available upon request. Sequencing data generated in this study have been deposited in the Sequence Raw Archive (https://www.ncbi.nlm.nih.gov/sra) under accession codes PRJNA950466; refer to for corresponding metadata and read statistics. The code for handling data from domain insertion library sequencing is available at: https://github.com/SavageLab/dipseq. The SPINE code is available at: https://github.com/schmidt-lab/SPINE. The version used for this study is archived at https://zenodo.org/badge/latestdoi/223953195. Processed read count data, along with R scripts to reproduce manuscript figures, is available at https://github.com/Schmidt-lab/AAV_Insertion_Profiling. A Shiny app to display 3D views of AAV insertion fitness maps in a web browser is available at https://github.com/Schmidt- <u>lab/shinyAAViewerR</u>.

## ACKNOWLEDGEMENTS

Some AAV productions used in this study were generated by the University of Minnesota Viral Vector and Cloning Core (Minneapolis, MN). We thank the University of Minnesota Genomics Center for providing NGS service and technical support. This work was also supported by the resources and staff at the University of Minnesota University Imaging Centers (UIC). The UIC RRID is SCR_020997. Transmission electron microscopy was carried out in the Characterization Facility, University of Minnesota, which receives partial support from the NSF through the MRSEC (Award Number DMR-2011401) and the NNCI (Award Number ECCS-2025124) programs.

## AUTHOR CONTRIBUTIONS

MDH, ACZ, and DS designed the study, with input from WRG and GA. MDH, ACZ., DN, and DS conducted the experiments with assistance from YH. MDH and DS analyzed the data and authored the manuscript. All authors have given approval to the final version of the manuscript.

---

**Supplemental Figure 1.**
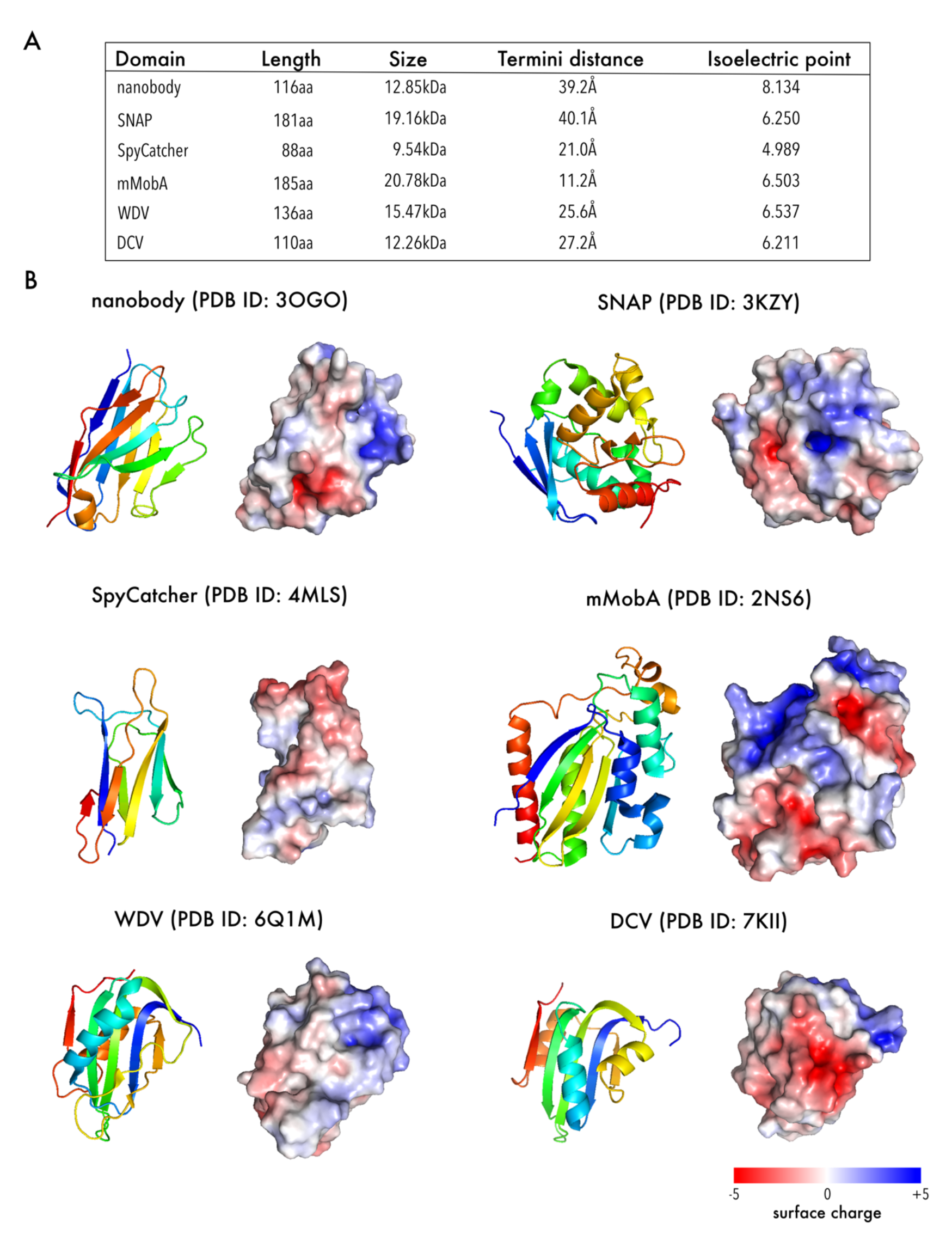
Properties of inserted domains. (A) Physical descriptions of domains used in this study. (B) Cartoon representation of domain structures (left) and surface representation with net surface charge shown as a gradient from red (negative), over white (neutral), to blue (positive).

**Supplemental Figure 2.**
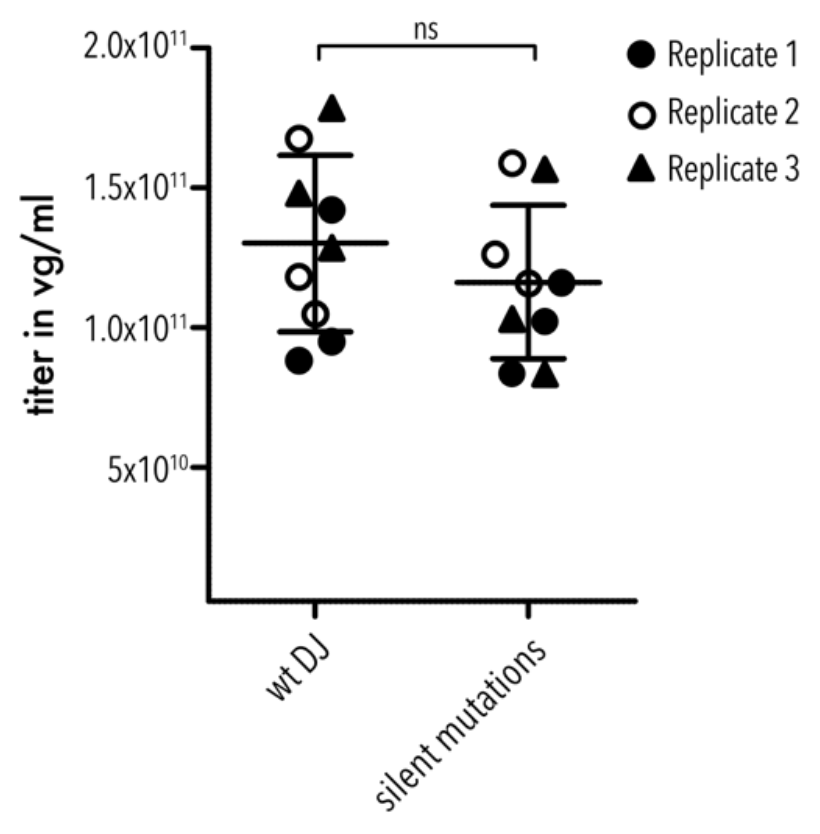
Packaging fitness of silent mutation variant compared to AAV-DJ. Crude lysate packaging titers quantified via qPCR. Three replicates with three technical replicates each were performed. Data are means ± SD. No statistical significance (ns) between AAV-DJ and the silent mutations variant by an unpaired, two-sided Student’s t-test (p-value: 0.3327).

**Supplemental Figure 3.**
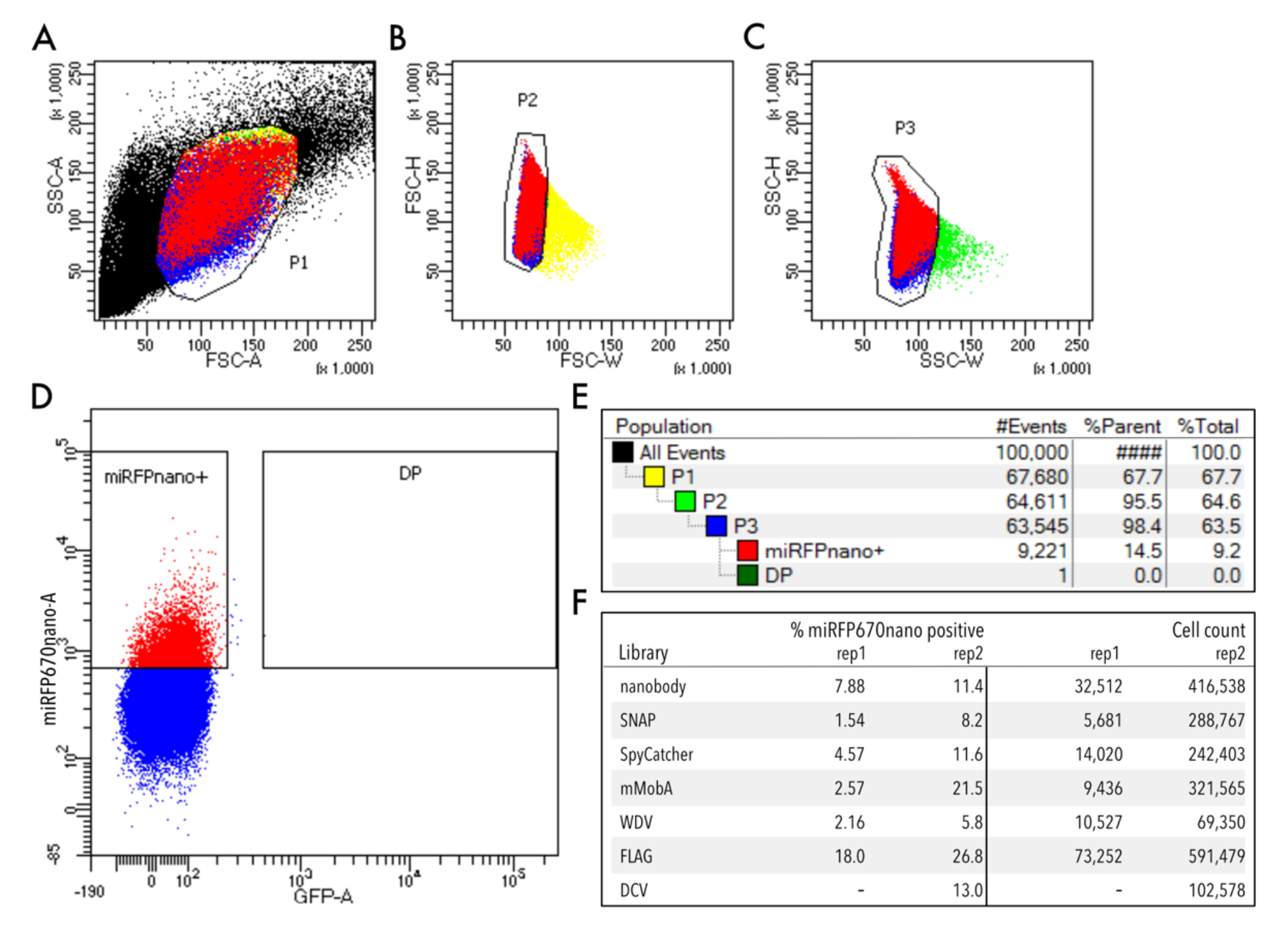
Infection fitness assay gating scheme. (A) Whole HEK293FT cells are gated on side (SSC-A) and forward scattering (FSC-A). (B-C) Forward scattering height (FSC-H), forward scattering width (FSC-W), and side scattering width (SSC-W) are used to gate single cells. (D) Cells are further gated using miRFP670nano as an infection marker (representative example). (E-F) Sort statistic for each gated cell population.

**Supplemental Figure 4.**
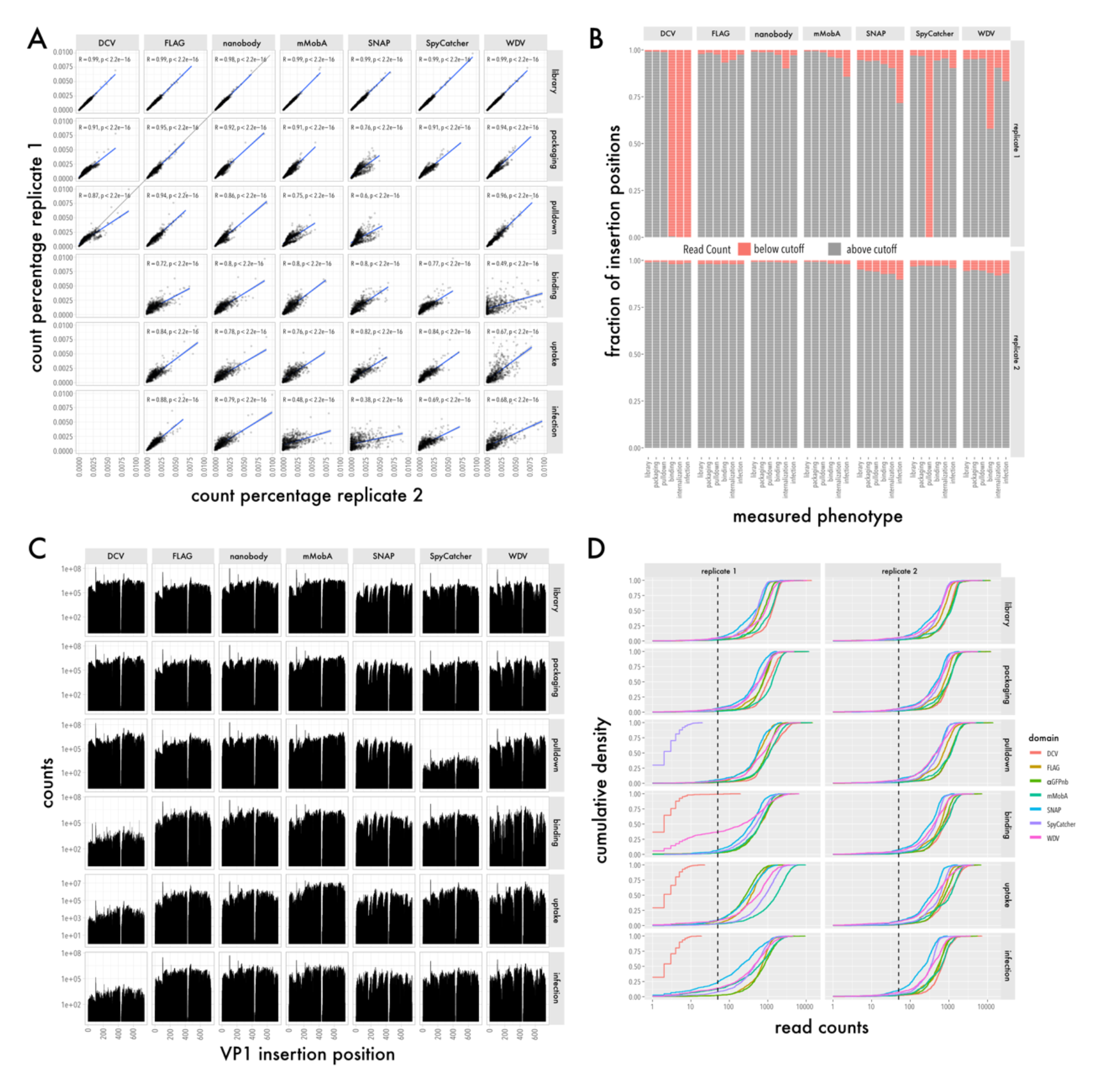
Fitness assay biological replicates, data completeness, and depth. (A) Read counts of replicate 1 are plotted against read counts of replicate 2 for the plasmid library, packaging, pulldown, binding, uptake, and infection assays. Data for all seven inserted domains are shown. Linear correlation was calculated (blue line). Pearson correlation coefficient and p values are shown. (B) Contingency plots showing the fraction of insertion positions below (red) and above (grey) the read count quantity cut-off of 50 reads for all seven domains and measured phenotypes. Data are shown for replicate 1 (top) and replicate 2 (bottom). (C) Total read counts of the plasmid library, packaging, pulldown, binding, uptake, and infection assays for each position are represented by insertion position. Data for all seven inserted domains are shown. (D) Cumulative density plots showing the read counts of the plasmid library, packaging, pulldown, binding, uptake, and infection assays for all seven inserted domains for replicate 1 and replicate 2. Cut off for sufficient read quantity was set to 50 reads and is represented by the black dashed lines.

**Supplemental Figure 5.**
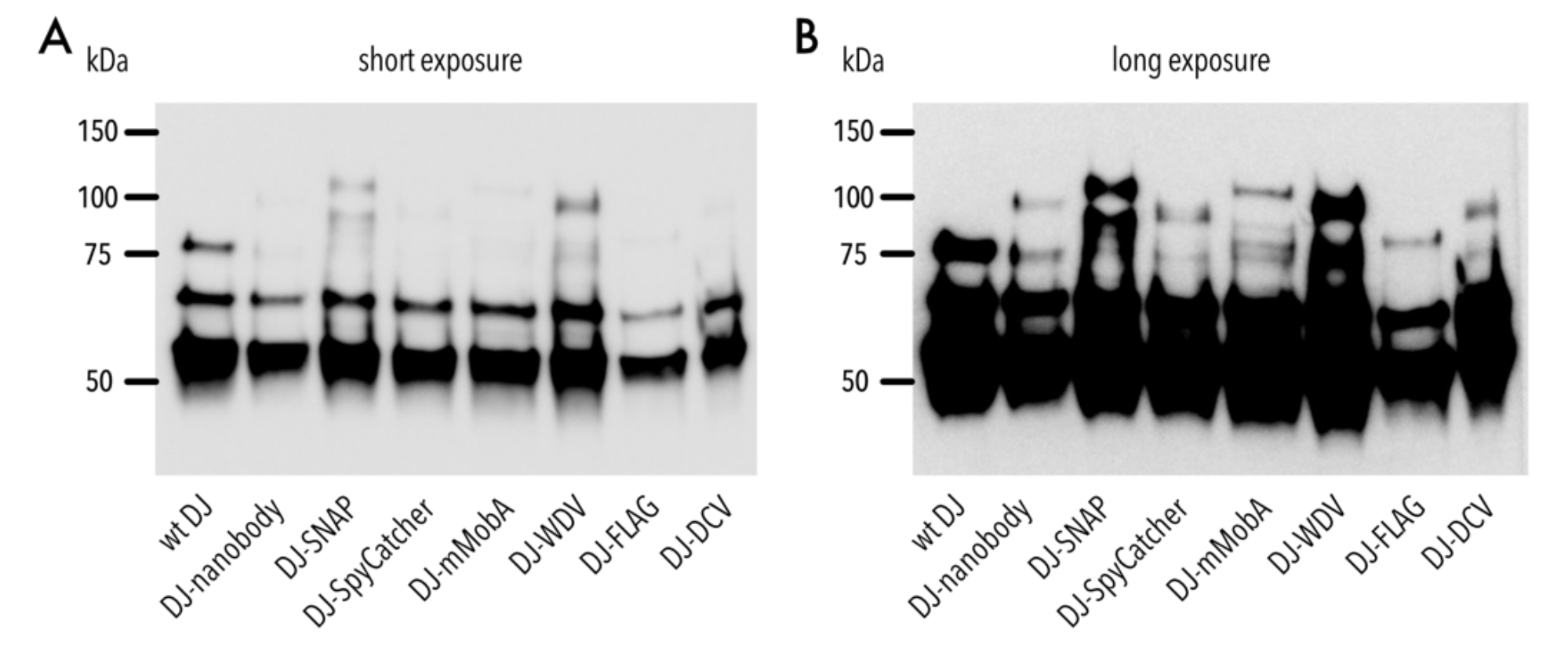
Western blot of AAV domain insertion libraries. (A-B) Representative Western blot image of AAV domain insertion libraries stained with B1 antibody (detecting VP1, VP2 and VP3 subunits) at a short exposure time (A) and long exposure time (B).

**Supplemental Figure 6.**
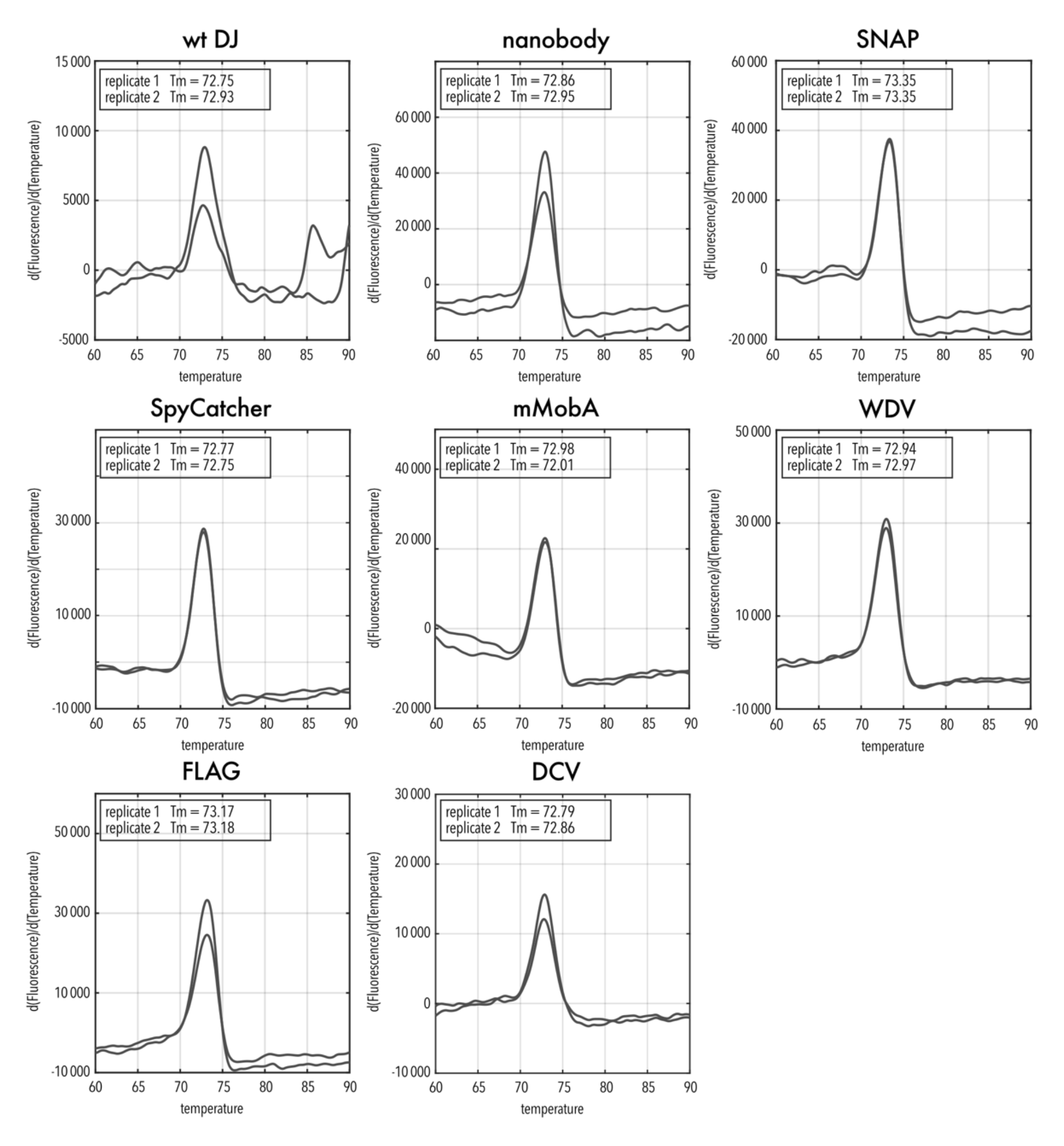
Thermal profiles of AAV domain insertion libraries. Data of two replicates are shown as "∂(Fluorescence)/∂(Temperature)" versus Temperature in °C.

**Supplemental Figure 7.**
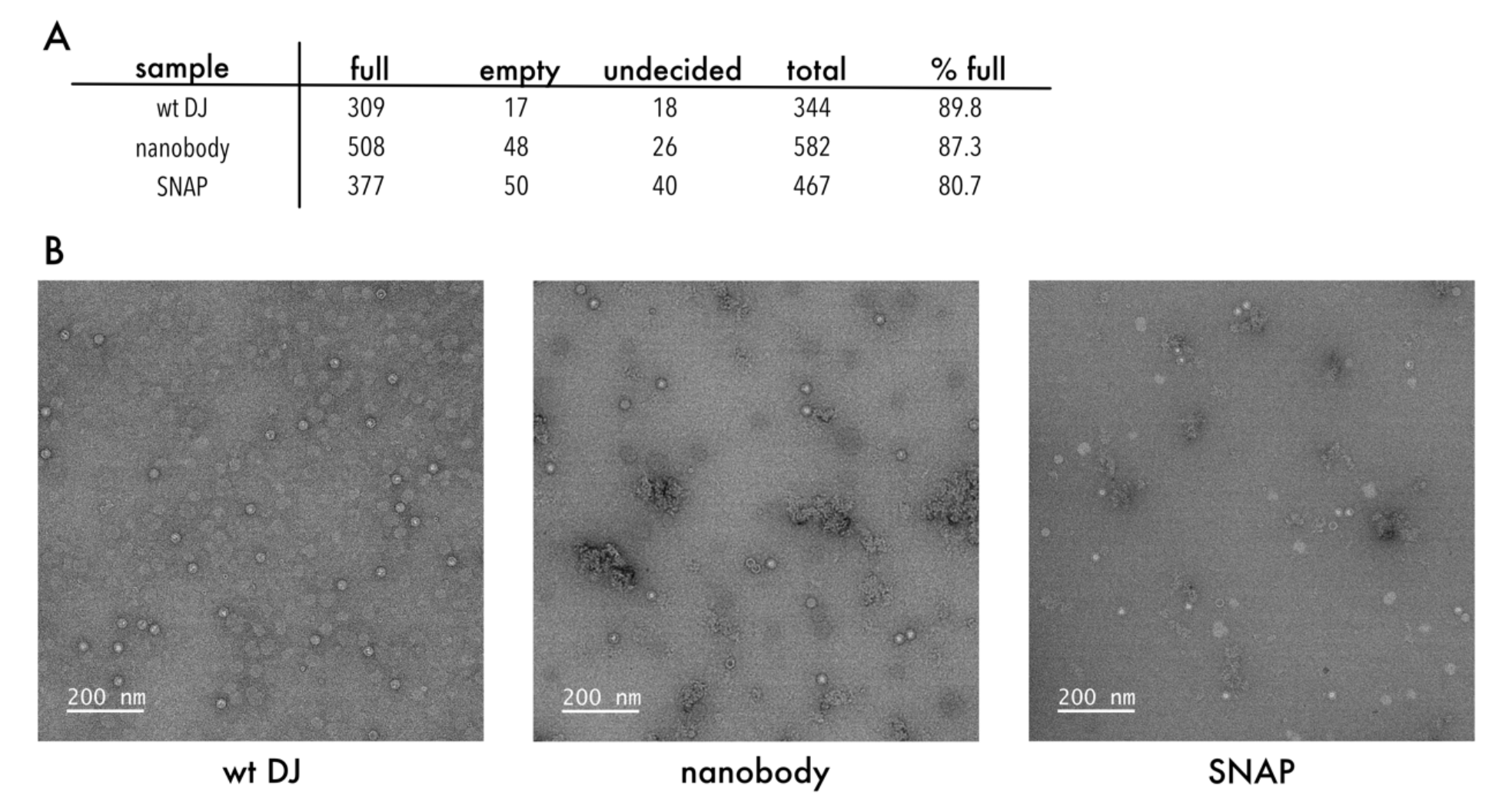
Quantification of empty to full capsid ratio by negative staining transmission electron microscopy. (A) More than 300 capsids were counted manually and grouped into full, empty, and undecided and the percentage of full capsids was calculated. (B) Representative transmission electron microscopy images of the indicated samples.

**Supplemental Figure 8.**
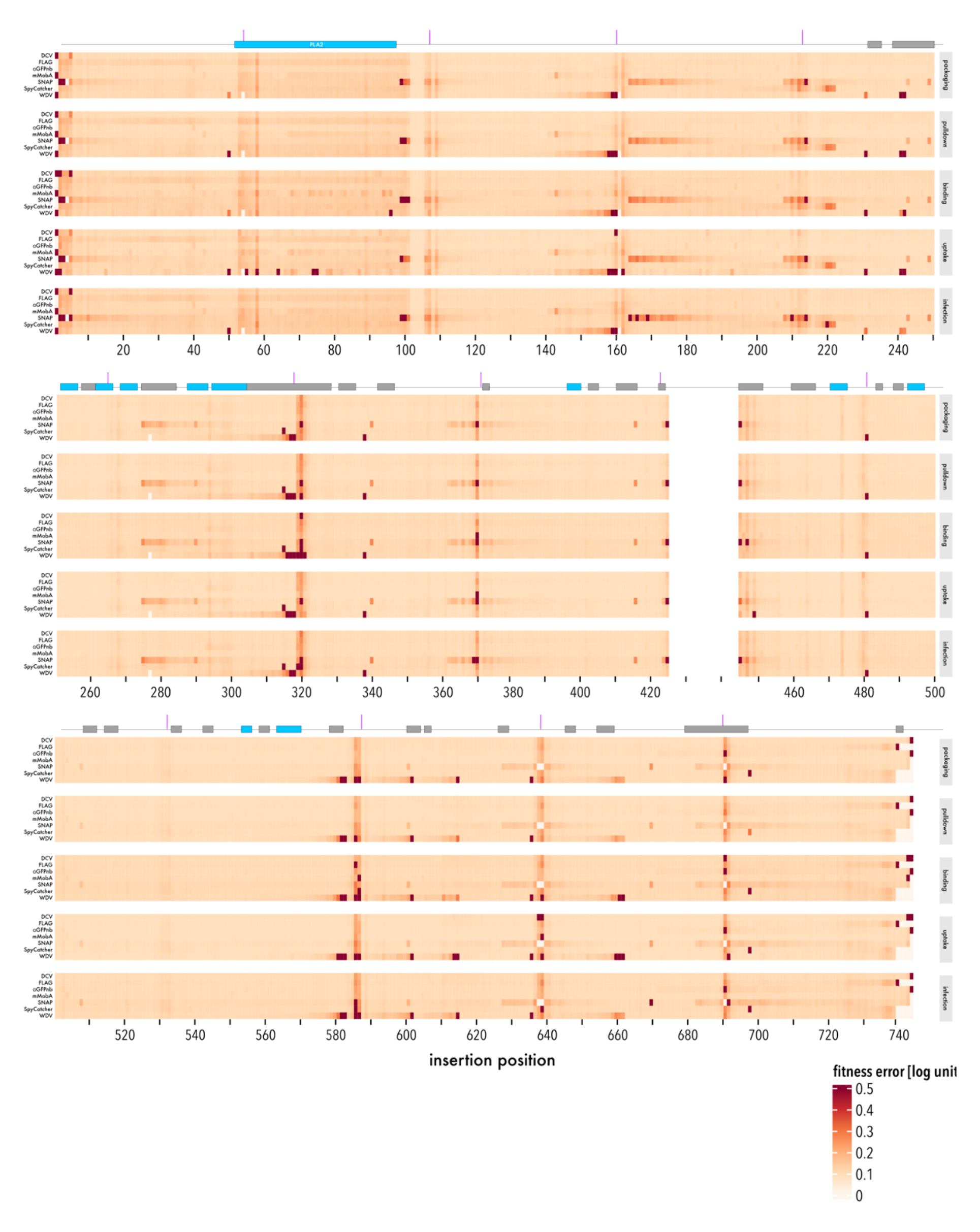
Accuracy of domain insertional profiling data. Standard errors of AAV fitness of all seven domain insertion libraries by insertion position and fitness assay. Secondary structure elements and VR1- 9 of the capsid are indicated on top. Boundaries of oligos from VP1 assembly using SPINE are indicated by purple vertical bars.

**Supplemental Figure 9.**
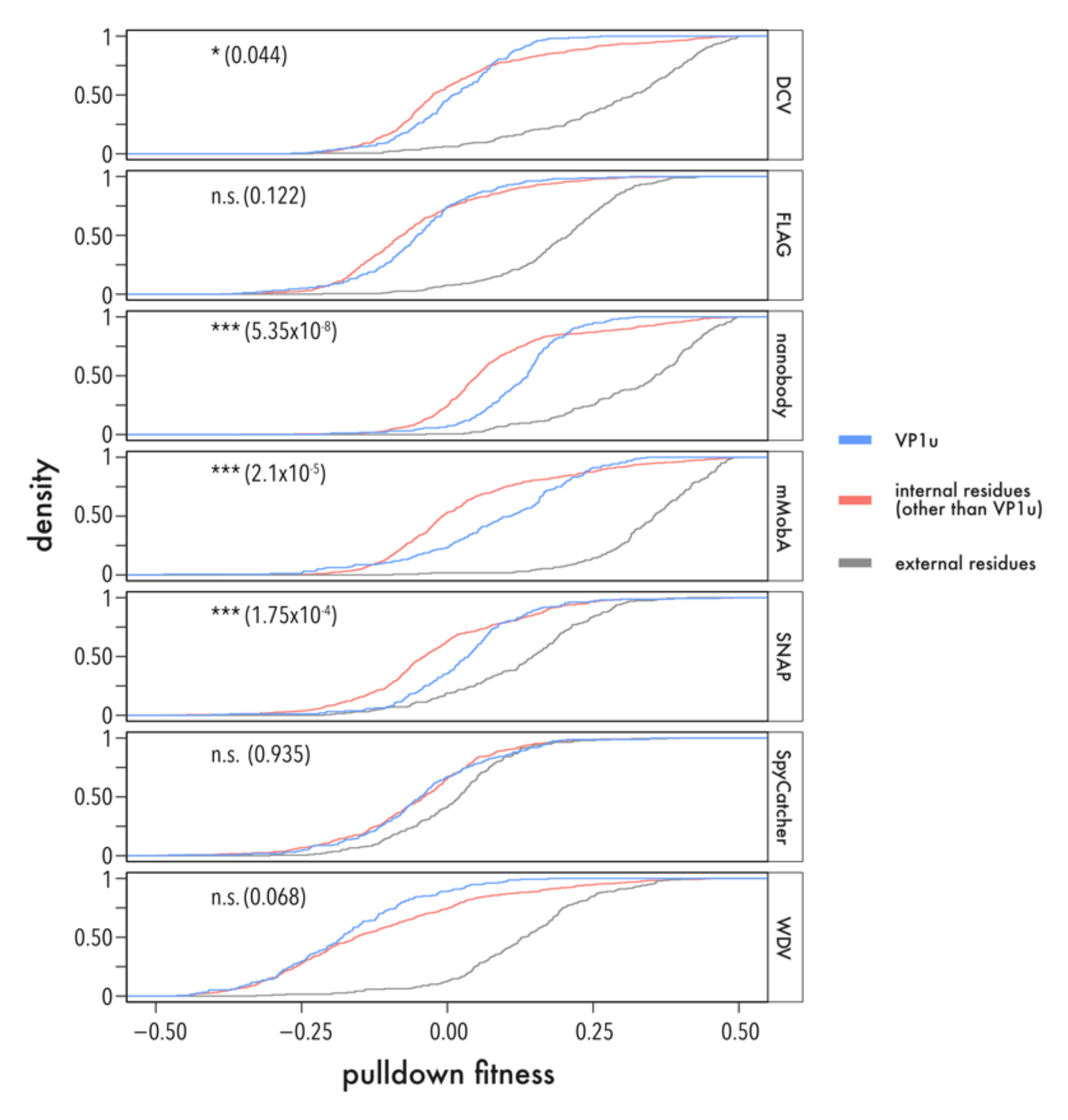
Distribution of pulldown fitness by residue location. Cumulative density function of pulldown fitness for each inserted motif stratified by residue location; within VP1u (blue), buried or exposed inside the capsid and not in VP1u (red), external residues (gray). Significance of distribution differences was tested using a two-sided, two-sample Kolmogorov-Smirnov test. Significance level and p values are shown.

**Supplemental Figure 10.**
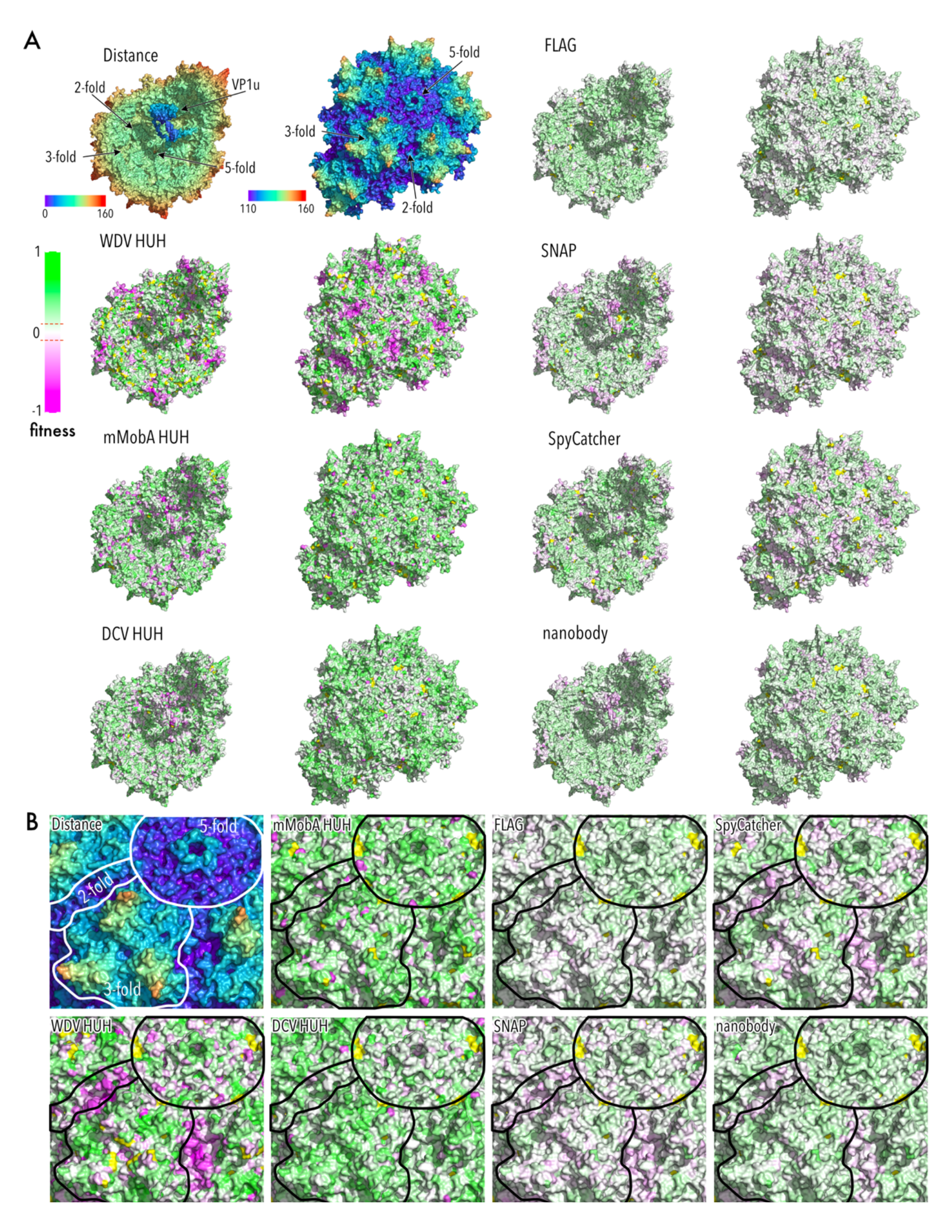
Binding fitness of AAV domain insertion libraries mapped to the capsid structure. (A) Top left corner: AAV-DJ capsid structure view from the inside (left) and outside (right) radially color-cued. 2-, 3-, and 5-fold axes are indicated. VP1u domain was modeled using RoseTTAFold (98) and manually positioned. All other structures show binding fitness heatmaps of the indicated domain insertions. Green indicates higher and magenta lower fitness than AAV-DJ (white) (RCSB PDB 7KFR). (B) Zoom of the outside structures from (A). 2-, 3-, and 5-fold axes are outlined.

**Supplemental Figure 11.**
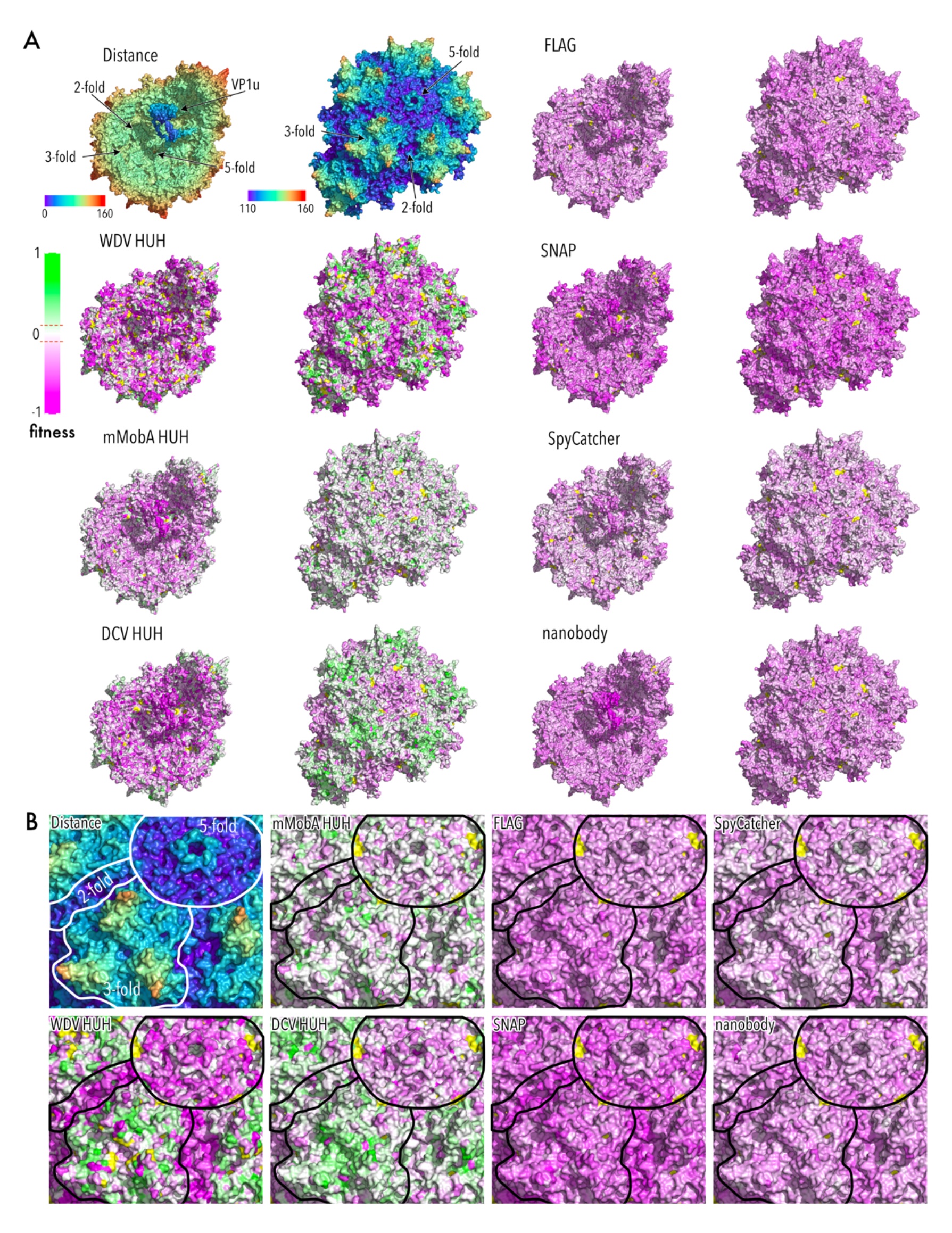
Uptake fitness of AAV domain insertion libraries mapped to the capsid structure. (A) Top left corner: AAV-DJ capsid structure view from the inside (left) and outside (right) radially color-cued. 2-, 3-, and 5-fold axes are indicated. VP1u domain was modeled using RoseTTAFold (98) and manually positioned. All other structures show uptake fitness heatmaps of the indicated domain insertions. Green indicates higher and magenta lower fitness than AAV-DJ (white) (RCSB PDB 7KFR). (B) Zoom of the outside structures from (A). 2-, 3-, and 5-fold axes are outlined.

**Supplemental Figure 12.**
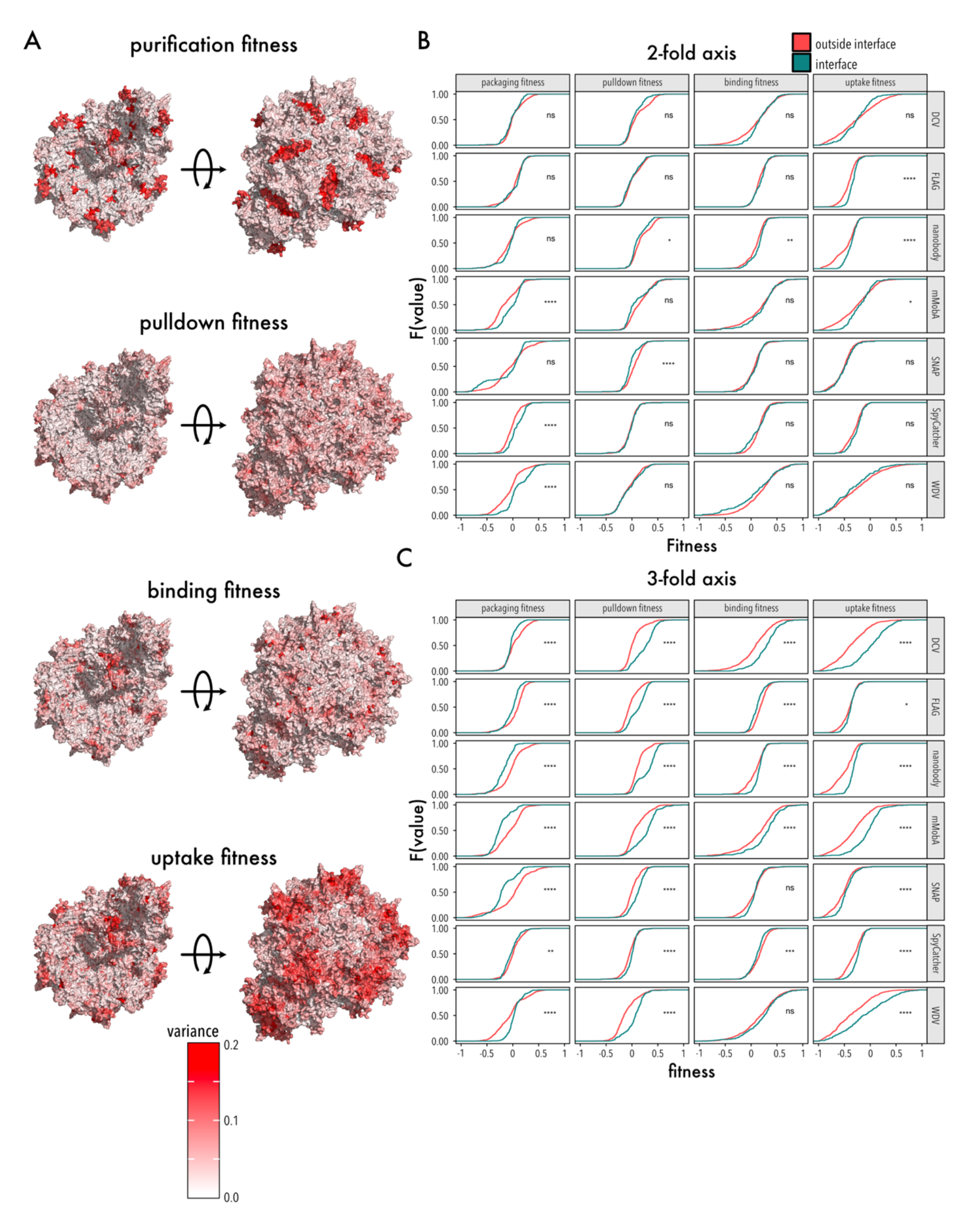
Variance of all measured fitness assays. (A) Variance of different fitness measures from all domain insertion libraries mapped to the AAV capsid structure. The capsid inside (left) and the capsid outside (right) are shown (RCSB PDB 7KFR). (B) Empirical cumulative density insertional fitness of residues within (petrol green) and outside (red) at the 2-fold axis (top) and 3-fold axis (bottom). Significance of distribution differences was tested using a two-sided, two-sample Kolmogorov-Smirnov test. Significance levels are shown (**** < 0.0001, *** < 0.001, ** < 0.01, * < 0.05).

**Supplemental Figure 13.**
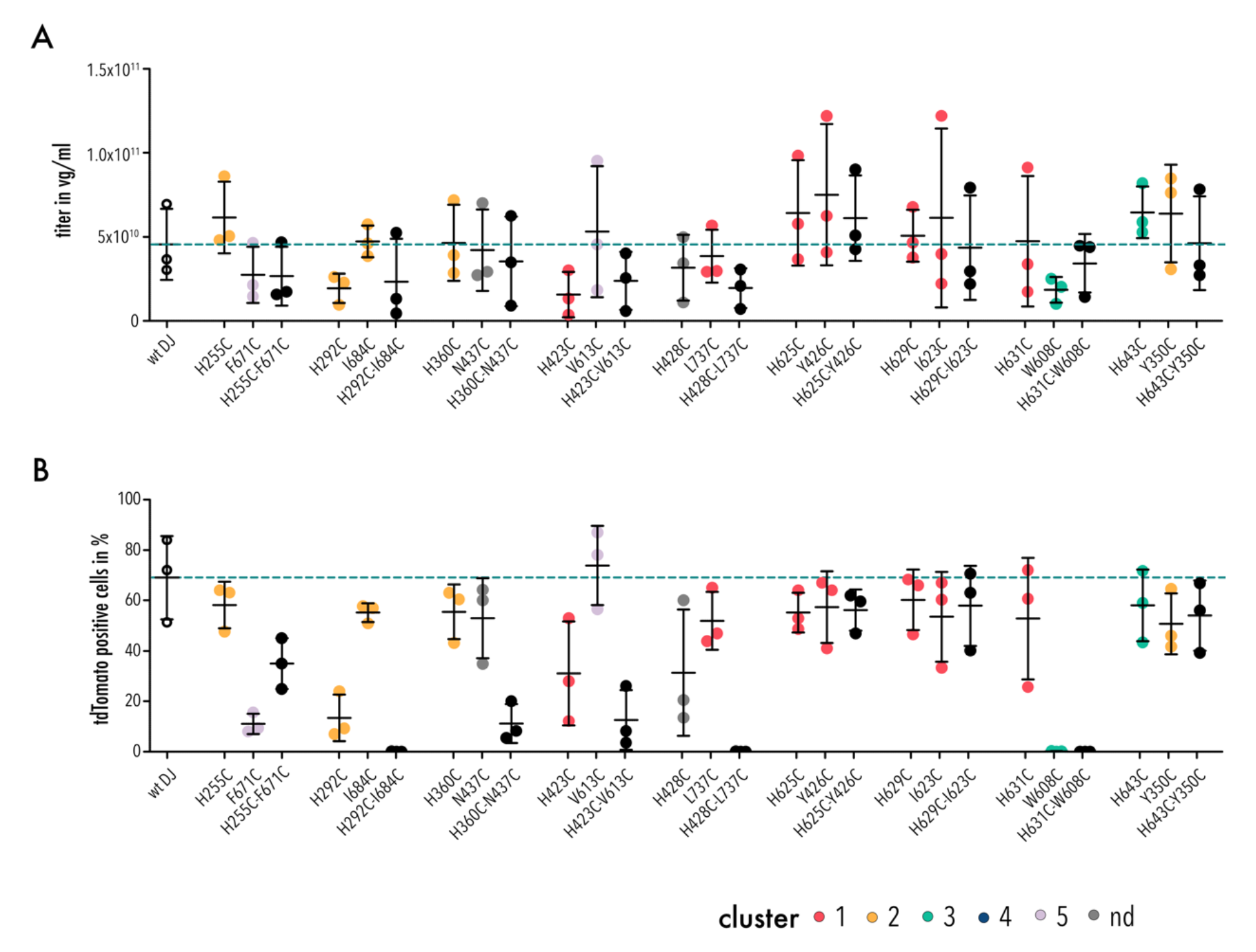
Packaging and infection fitness of cysteine mutants. (A) Crude lysate packaging titers quantified via qPCR. (B) Infection fitness quantified by measuring the percentage of tdTomato positive cells 48 hours post transduction with an MOI of 1E4 vg. Data are means ± SD. Data points of single mutants are colored by cluster membership, missing residues are gray, wildtype AAV-DJ fitness is shown as open circles and horizontal dashed line.

**Supplemental Figure 14.**
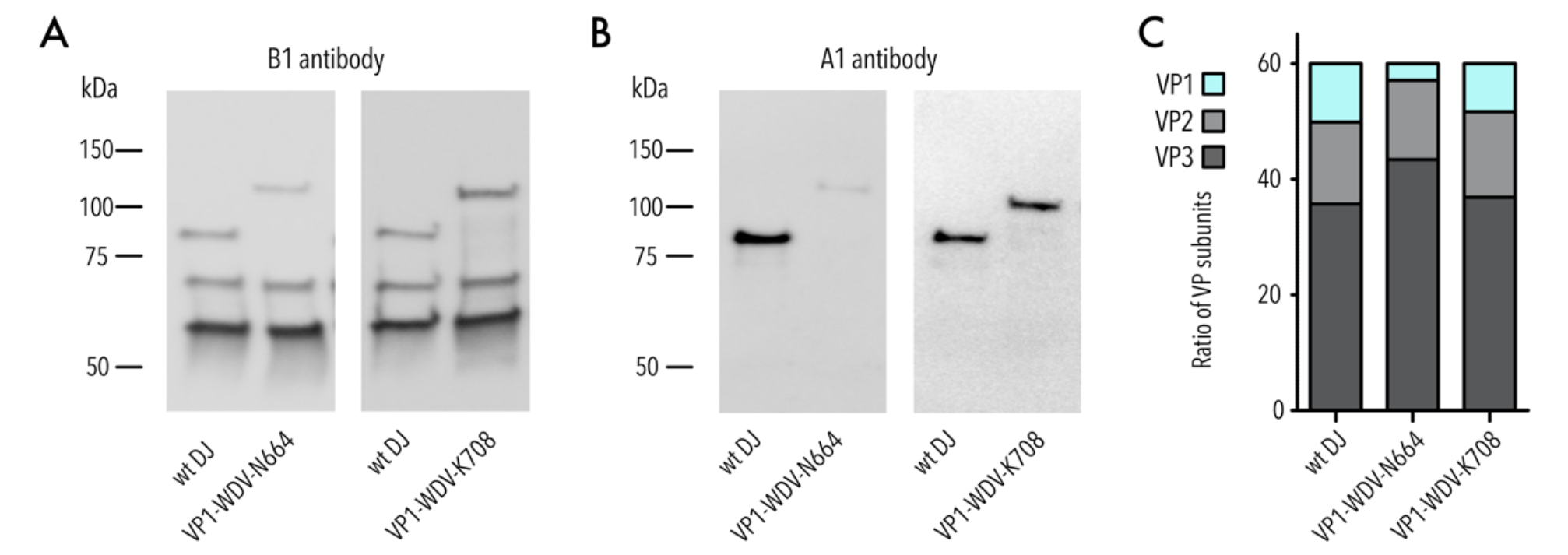
Quantification of VP ratios of WDV insertion variants N664 and K708. (A) Representative Western blot image of AAV domain insertion libraries stained with B1 antibody (detecting VP1, VP2 and VP3 subunits). (B) Representative Western blot image of AAV domain insertion libraries stained with A1 antibody (detecting VP1 subunits). (C) Western blot quantification of VP1, VP2, and VP3 subunits. Data are means (n=3).

**Supplemental Table 1.**
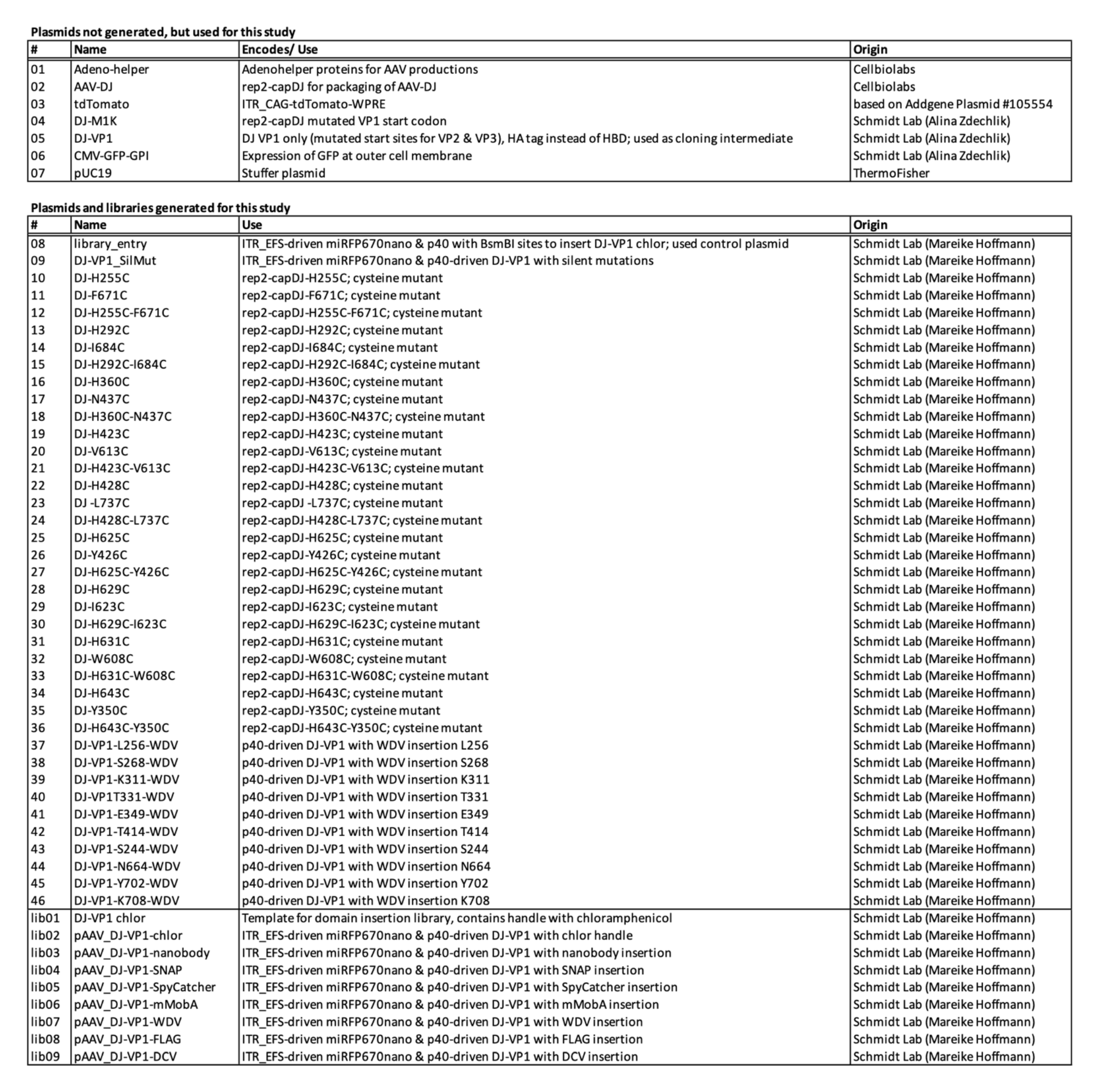
Plasmids and libraries used in this study.

**Supplemental Table 2.**
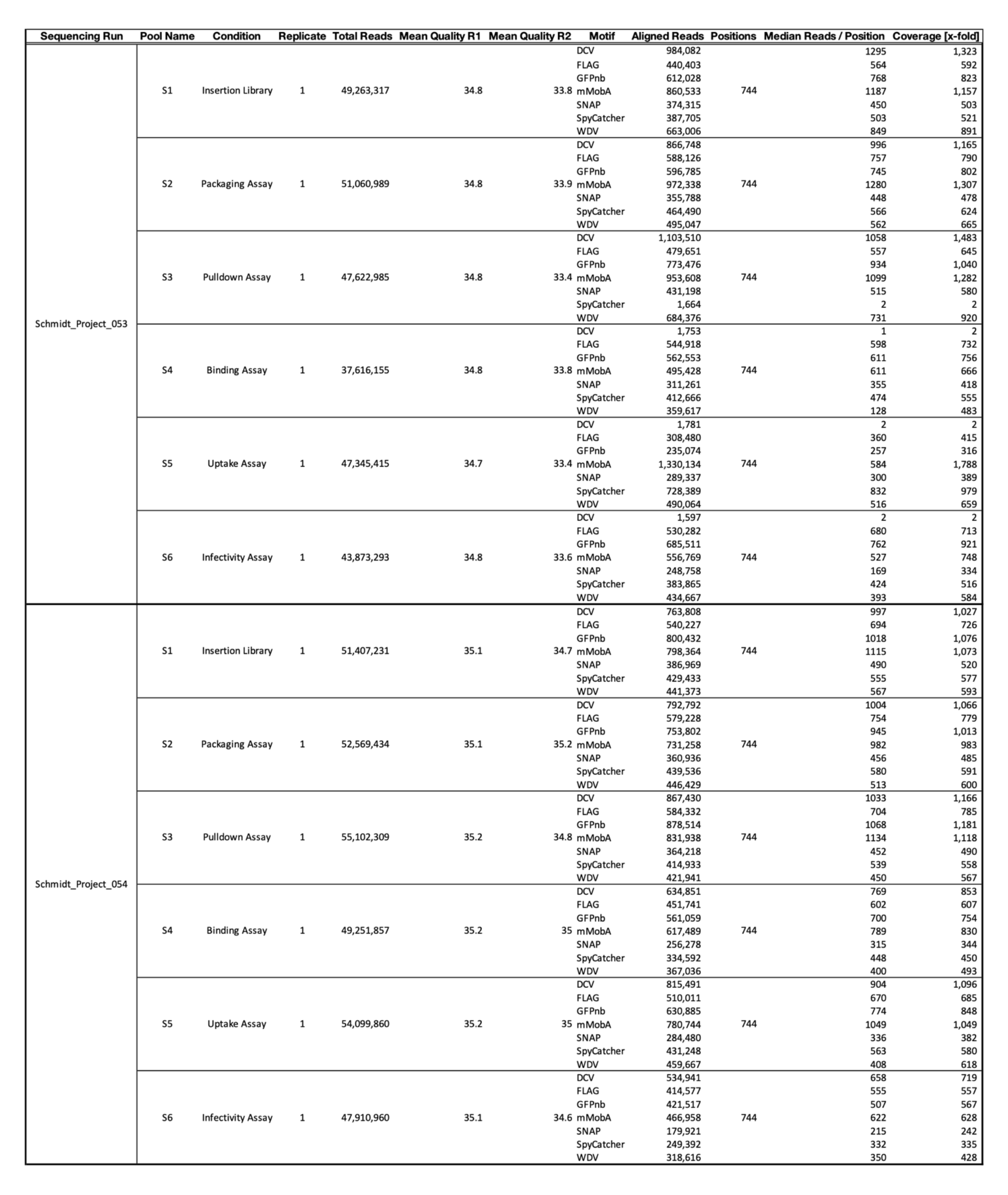
Sequencing statistics.

**Supplemental Table 3.**
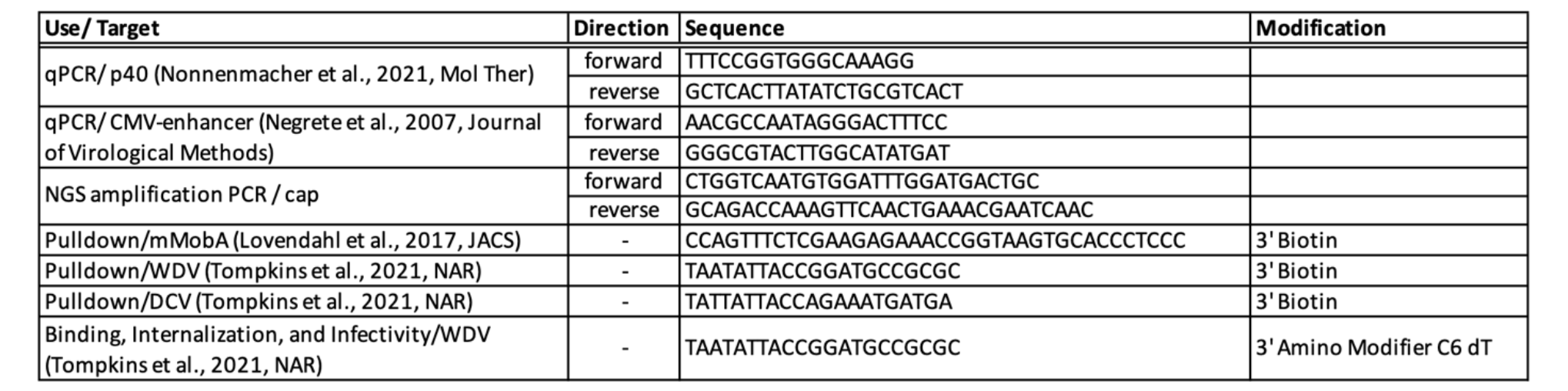
DNA oligos used in this study.

